# Sparse modeling of interactions enables fast detection of genome-wide epistasis in biobank-scale studies

**DOI:** 10.1101/2025.01.11.632557

**Authors:** Julian Stamp, Samuel Pattillo Smith, Daniel Weinreich, Lorin Crawford

## Abstract

The lack of computational methods capable of detecting epistasis in biobanks has led to uncertainty about the role of non-additive genetic effects on complex trait variation. The marginal epistasis framework is a powerful approach because it estimates the likelihood of a SNP being involved in any interaction, thereby reducing the multiple testing burden. Current implementations of this approach have failed to scale to large human studies. To address this, we present the sparse marginal epistasis (SME) test, which concentrates the scans for epistasis to regions of the genome that have known functional enrichment for a trait of interest. By leveraging the sparse nature of this modeling setup, we develop a novel statistical algorithm that allows SME to run 10 to 90 times faster than state-of-the-art epistatic mapping methods. In a study of blood traits measured in 349,411 individuals from the UK Biobank, we show that reducing searches of epistasis to variants in accessible chromatin regions facilitates the identification of genetic interactions associated with regulatory genomic elements.

## Introduction

Genome-wide association studies (GWAS) have identified thousands of genetic loci linked with various complex traits and common diseases, offering valuable insights into the genetic foundations of phenotypic variation [1]. As of late, there have been many efforts to estimate proportions of genetic variance beyond what is attributable to additive effects [2–6]. Epistasis, which refers to interactions between genetic loci, is thought to play a key role in the constituting the genetic basis of evolution [7, 8]. While many studies have shown epistasis to be pervasive in model organisms [9–11], controversies remain with respect to its role in humans [12]. For example, some epistatic interactions identified in association mapping studies can be explained by additive effects of unobserved variants [13]. Though previous studies have shown that genetic variance is mainly additive [9, 14], these conclusions have recently been challenged [5].

Numerous statistical methods have been developed to identify single nucleotide polymorphisms (SNPs) that contribute to epistasis. Traditional approaches focus on explicitly detecting significant interactions through exhaustive or probabilistic searches utilizing frequentist tests, Bayesian inference, and machine learning techniques [15–18]. With advancements in sequencing technologies, many contemporary genome-wide association studies (GWAS) are conducted on biobank-scale data sets, comprising hundreds of thousands of individuals genotyped at millions of markers and phenotyped for thousands of traits [1, 19, 20]. This is crucial as the effect of epistatic interactions is hypothesized to be small for many traits [12, 14], and traditional search algorithms are known to be most powered when large training data sets are available [14, 15]. However, despite efficient computational improvements, exploring large combinatorial domains continues to pose a challenge for epistatic mapping studies. With a lack of a priori knowledge about which epistatic loci to prioritize, exploring all possible combinations of genetic variants can result in low statistical power after correcting for multiple hypothesis tests (e.g., there are *J* choose 2 possible pairwise combinations for a study with *J* SNPs).

As an alternative to traditional exhaustive search methods, the marginal epistasis framework was developed to estimate the combined pairwise interaction effects between a focal SNP and all other variants in the data set. The “Marginal Epistasis Test” (MAPIT) evaluates each SNP individually and identifies candidates involved in epistasis without requiring the identification of their exact interacting partners [2]. Recently, the concept of marginal epistasis has been leveraged to estimate the contribution of non-additive heritability in complex traits using GWAS summary statistics [5]. It has also been extended to explore the importance of gene-by-environment interactions on complex traits [21]. Theoretically, MAPIT is formulated as a linear mixed model where the random effects and corresponding variance components are estimated using a method-of-moments (MoM) algorithm [22, 23]. Although MAPIT mitigates the reduction of power due to multiple testing burden, its implementation on data sets with large sample sizes remains computationally intensive [2]. Specifically, the computational complexity scales linearly with the number of SNPs and at best quadratically with the number of individuals, making it suitable for moderately sized GWAS applications but infeasible for biobank-scale studies[24, 25]. Efforts have been made to address this limitation, such as the “fast marginal epistasis test” (FAME) [4], which leverages a stochastic MoM framework and introduces both computationally efficient stochastic trace estimators [26] and innovative methods to expedite matrix multiplication [27]. However, despite these advancements, further work is necessary to scale the method to genome-wide applications.

This work introduces the sparse marginal epistasis (SME) test, which focuses on searching for epistasis in regions of the genome with known functional enrichment [28] related to a trait of interest. This method has two main advantages. First, it prioritizes candidate regions likely to involve epistatic gene action. Studies have indicated that variants in coding regions account for less than 10% of the phenotypic variance in many traits and diseases [29]. Consequently, the remaining heritability is attributed to regions expected to play a regulatory role [28–30] and that are active in trait-specific tissue [31, 32]. Second, the sparse nature of this approach leads to more efficient estimators [23, 33] for model parameters and allows SME to operate significantly faster than existing methods such as MAPIT and FAME. Through detailed simulations, SME demonstrates effective type I error control and improved power compared to previous methods. Furthermore, utilizing information from DNase I hypersensitivity sites in ex vivo human erythroid differentiation [34], we use SME to analyze hematology traits in individuals from the UK Biobank [19] and identify genetic interactions associated with regulatory genomic elements.

## Results

### SME overview

The sparse marginal epistasis (SME) test performs a genome-wide search for SNPs involved in genetic interactions while conditioning on information derived from functional genomic data (Fig. 1a). Assume that we are analyzing a GWAS data set where **X** is an *N* × *J* matrix of genotypes with *J* denoting the number of SNPs (each of which is encoded as 0, 1, 2 copies of a reference allele at each locus *j*) and **y** is an *N*-dimensional vector of measurements of a quantitative trait. Also assume that we have access to an external reference *S* that encodes some additional biological information about the trait being studied. By examining one SNP at a time (indexed by *j*), SME fits the following linear mixed model (Methods)

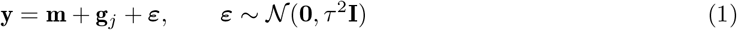

where **m** =∑_*l*_ **x**_*l*_ *β*_*l*_ is the combined additive effects from all SNPs and **g**_*j*_ = ∑_*l≠j*_ (**x**_*j*_ ○ **x**_*l*_) *α*_*l*_ · 𝟙_𝒮_(*w*_*l*_) represents the effects of a subset of pairwise interactions involving the *J*-th SNP. The key to this formulation is that the inclusion of the interaction between the *J*-th and *l*-th SNPs is based on an indicator function

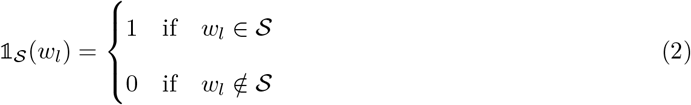

where *w*_*l*_ encodes information about the *l*-th SNP. For example, if testing for epistatic effects in red blood cell traits, we can incorporate information about regulatory regions during erythroid differentiation into the test (Fig. 1b). In this case, 𝒮 could be a set of genomic regions for which DNase-seq implicates chromatin accessibility and *w*_*l*_ could encode the physical location of the *l*-th SNP on the genome. Here, 𝟙_𝒮_(*w*_*l*_) = 1 if the *l*-th SNP is located in one of these regions (i.e., *w*_*l*_ ∈ 𝒮). This means that, while all SNPs are tested for marginal epistasis, only their interactions with SNPs included in Eq. (2) are considered.

**Figure 1.**
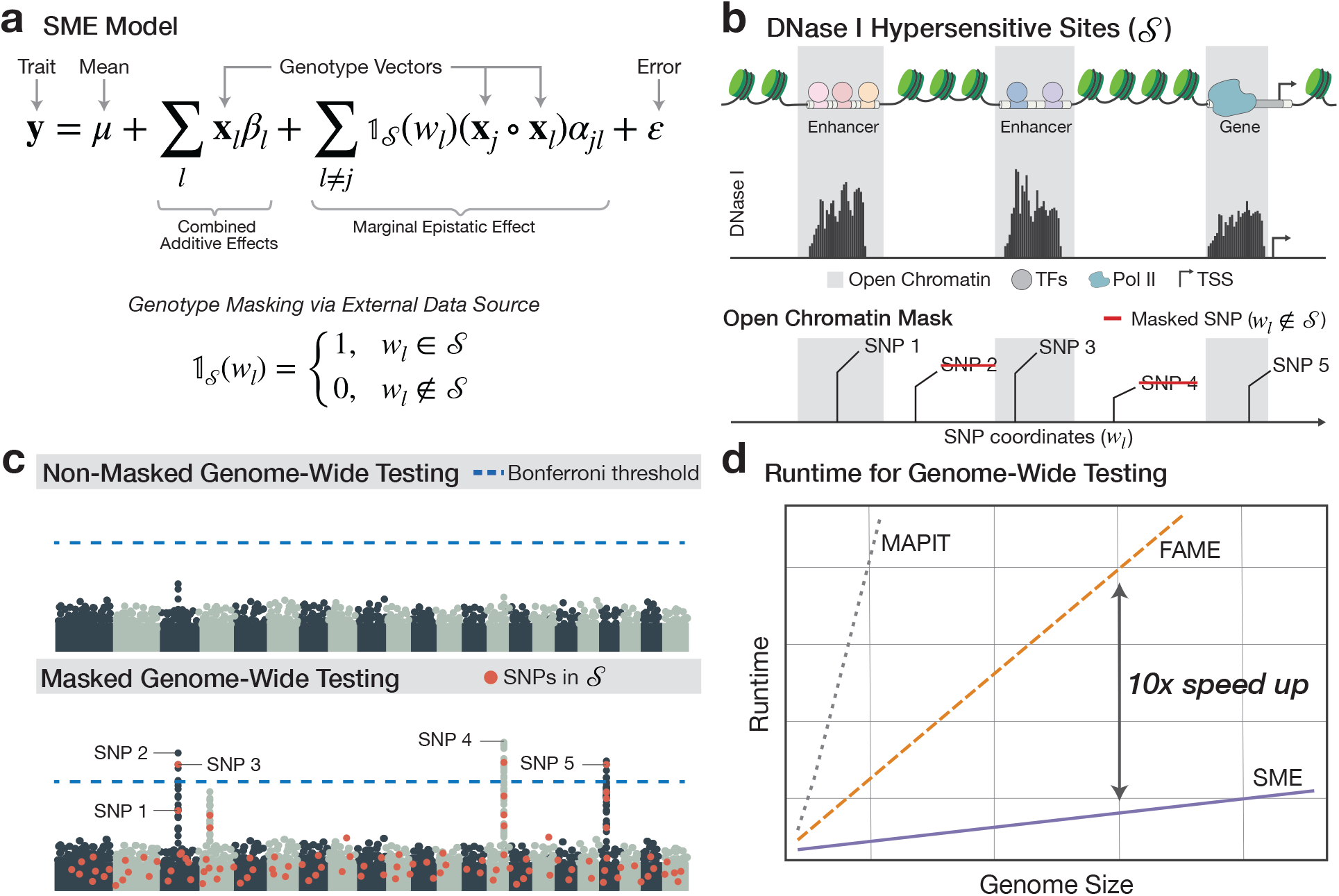
Schematic overview of the sparse marginal epistasis (SME) test. **(a)** SME examines one SNP at a time and estimates marginal epistatic effects—the combined pairwise interaction effects between a *j*-th focal SNP and other variants on the genome (indexed by *l* ≠ *j*). The key to SME is that it incorporates genomic data 𝒮 through a binary indicator function 𝟙_𝒮_ (*w*_*l*_) where *w*_*l*_ provides information about the *l*-th background SNP. This creates a mask to only search for interactions in regions of the genome with known functional enrichment related to a trait of interest. **(b)** As an example, let data on DNase I hypersensitive sites (DHS) be used for 𝒮. In this case, SME restricts the marginal epistasis test to assessing interactions between each focal SNP and variants with genomic coordinates that fall within open chromatin and regulatory regions. The DNase-seq signal is converted into a binary mask, excluding variants located in regions with closed chromatin (i.e., variants with coordinates *w*_*l*_ ∉ 𝒮). **(c)** SME tests every SNP genome-wide. Masking results in improved power to detect marginally epistatic variants versus the traditional non-masking approach. **(d)** SME uses computationally efficient estimators to enable genome-wide testing on biobank-scale data sets. It achieves runtimes 10 × to 90× faster than current state-of-the-art methods.

The benefits of this approach are twofold. First, by limiting the search to regions of the genome that are most likely to be functionally associated with a phenotype, SME produces significantly more efficient estimators which leads to an increase in power (Fig. 1c). Second, by masking out sets of variants for each test on a focal SNP, SME leverages a novel MoM algorithm to substantially improve its scalability for genome-wide analyses (Fig. 1d). Specifically, SME introduces an approximation to the efficient stochastic trace estimator [4, 24, 26, 35] which allows the algorithm to avoid repeating costly matrix computations when estimating model parameters across each SNP that is being tested (Figs. S1-S3).

Probabilistically, SME assumes that **m** ∼ 𝒩 (**0**, *ω*^2^**K**) where the covariance matrix **K** = **XX**^**⊤**^ /*J* accounts for the relatedness between individuals in the data and the corresponding component *ω*^2^ models the phenotypic variance explained by additive effects. The second term can be written as **g**_*j*_ ∼ 𝒩 (**0**, *σ*^2^**G**_*j*_) where the covariance matrix **G**_*j*_ represents all pairwise interactions involving the *j*-th SNP that have not been masked out according to the set of indicator functions {𝟙_*S*_(*w*_*l*_)}_*l* ≠ *j*_. The key to SME is that the variance component *σ*^2^ measures SNP-specific contribution to the non-additive genetic variance. Therefore, to identify significant nonzero marginal epistatic effects, the model assesses the null hypothesis *H*_0_ : *σ*^2^ ≤ 0 for each SNP in the data set. After SME estimates model parameters, it uses a calibrated one-sided z-score (i.e., normal) test to derive *P*-values.

### SME scales to biobank genome-wide association studies

To compare the expected central processing unit (CPU) computation for conducting a biobank-scale genome-wide analysis using SME, FAME [4], and MAPIT [2], we measured the average runtime per SNP for each method on an Intel(R) Xeon(R) Platinum 8268 CPU using a single core (i.e., no parallel processing). Here, we used genotype data from 349,411 individuals of self-identified European ancestry in the UK Biobank with 543,813 SNPs after quality control (Methods). The memory requirements for MAPIT are prohibitively high, requiring resources on the order of terabytes for biobank-scale data sets with hundreds of thousands of observations.

We find that SME performs genome-wide testing 10 × faster than FAME and 90× faster than MAPIT (Fig. 2). While analyzing the complete data set with a single core, SME requires only 3.7 days of runtime compared to FAME and MAPIT requiring 38.4 and 324 days, respectively. The greatest speedup in SME is achieved by approximating the stochastic trace when estimating model parameters (Fig. S1). This allows for computations involving large genetic relatedness matrices to be reused across multiple tests for different focal SNPs. Even with the stochastic trace approximations, performing matrix calculations at biobank-scale still takes minutes. However, the ability to share these computations across multiple variants significantly reduces the overall computational burden (Figs. S2 and S3). The proposed approach enables SME to be effectively applied to the UK Biobank, facilitating genome-wide association studies of epistasis.

**Figure 2.**
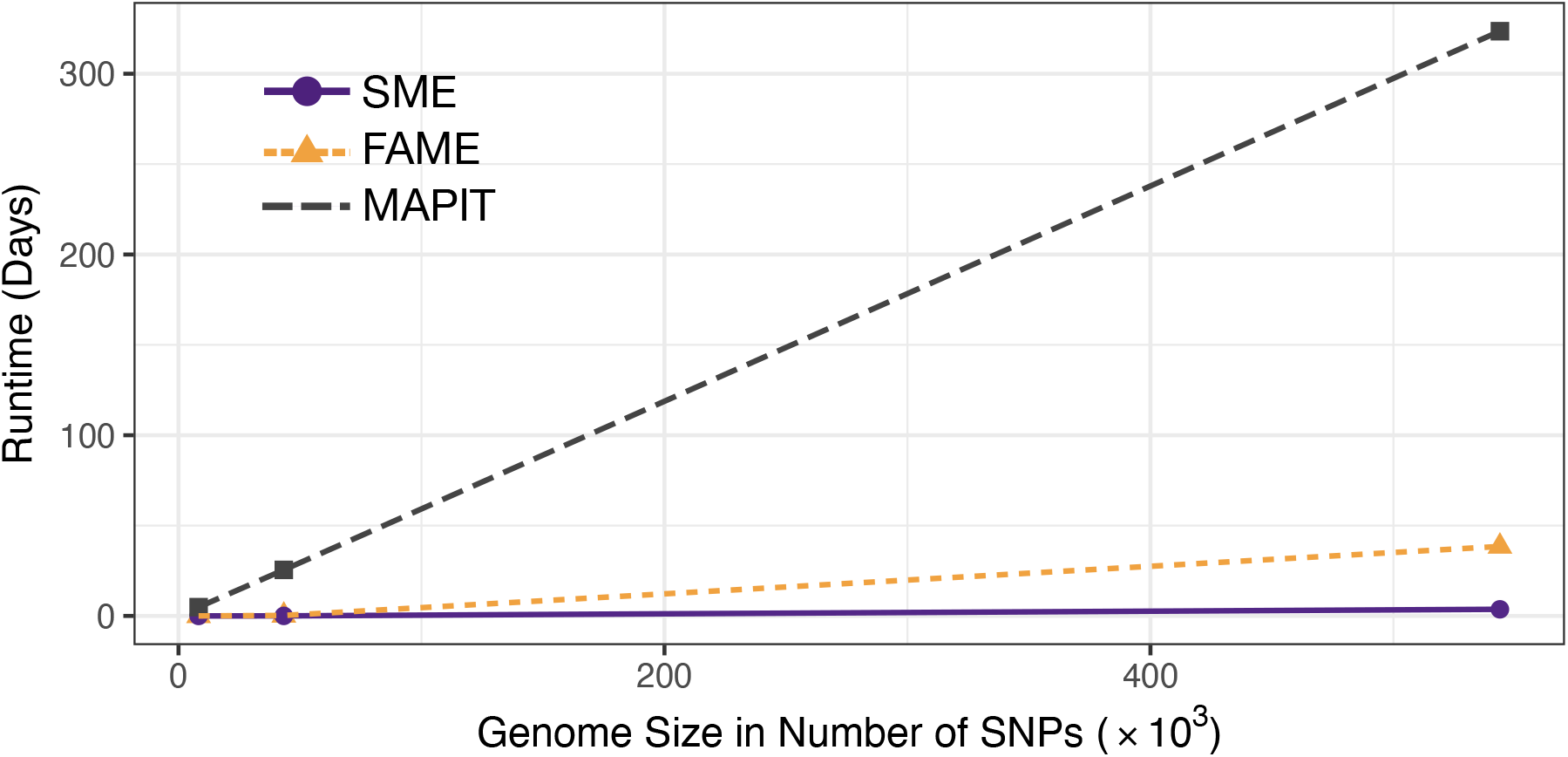
Computational time for running SME and other marginal epistatic approaches on biobank-scale data as a function of the genome size. The other methods compared include FAME [4] and MAPIT [2]. Here, we analyze genotype data from a fixed set of 349,411 individuals from the UK Biobank and vary the genome size. All results were computed on a single core of an Intel(R) Xeon(R) Platinum 8268 central processing unit (CPU). Total runtime was calculated based on the average runtime per SNP and parallel processing on a cluster with 960 CPUs available. Both SME and FAME were set to have the same hyperparameter configurations (e.g., the number of random vectors). SME also used a binary mask that contained 5,000 unmasked SNPs, and its stochastic trace approximation was applied such that sets of 250 focal SNPs shared the same random vectors. MAPIT could not be directly compared due to its excessive memory requirements for data sets of this size. Instead, the runtime for MAPIT was measured on smaller sample sizes and extrapolated to a sample size of 349,411. This extrapolation assumed quadratic scaling with the number of individuals and linear scaling with the number of SNPs.

### SME is a well-calibrated test and conserves type I error rates

We generate synthetic phenotypes using a linear model with real genotypes from chromosome 1 of white British individuals in the UK Biobank [2, 3, 5]. After quality control, we had a data set of 349,411 individuals and 43,332 SNPs (Methods). Under the null model, we simulate traits consisting of only additive effects. Here, we randomly sample 10% of the SNPs and scale their effect sizes such that they explain 40% of the total phenotypic variance.

We simulate external data sources (𝒮) to be used when generating a mask for the marginal epistatic covariance matrix **G**_*j*_ in SME. Recall that these external data sources are intended to give alternative insight into the importance of SNPs and are used to induce sparsity in the modeled gene-interactions by dropping interactions with “unimportant” variants. We consider two scenarios in our simulations (Fig. S4). In the first, SNPs deemed important in the external data source are sparsely sampled with uniform probability from all variants. As a result the modeled gene-interactions are evenly distributed along the chromosome. We will refer to this scenario as inducing *uniform sparsity* in the SME model. In the second scenario, we randomly sample one central seed SNP and define variants in a block around it as important. We will refer to this scenario as one that induces *localized sparsity*. In these simulation experiments, we assess the calibration of SME using both types of external data sources as a function of the percentage of total variants that are masked (varying between 0%, 95%, and 99%) and the number of individuals being analyzed (varying between 20k, 50k, 100k and 300k randomly subsampled individuals).

Under the null model, we find that SME produces well-calibrated *P*-values and unbiased variance component estimates (Fig. 3) using a uniformly sparse mask. Specifically, higher levels of sparsity lead to more accurate estimates of the marginal epistatic variance components. We also see precision in its estimates improve as the sample size increases, which is expected since SME uses a normal test to compute *P*-values for each SNP. Overall, this translates to SME preserving empirical type I error rates estimated at significance levels *α* = 0.05, 0.01, and 0.001, respectively (Table 1). In contrast, FAME produces inflated test statistics as the number of samples in a data set grows. Note that we do not include a comparison with MAPIT here due to its inability to scale to biobank settings (see Crawford *et al*. [2] for assessment of its calibration on small-to-moderately sized data).

**Table 1.**
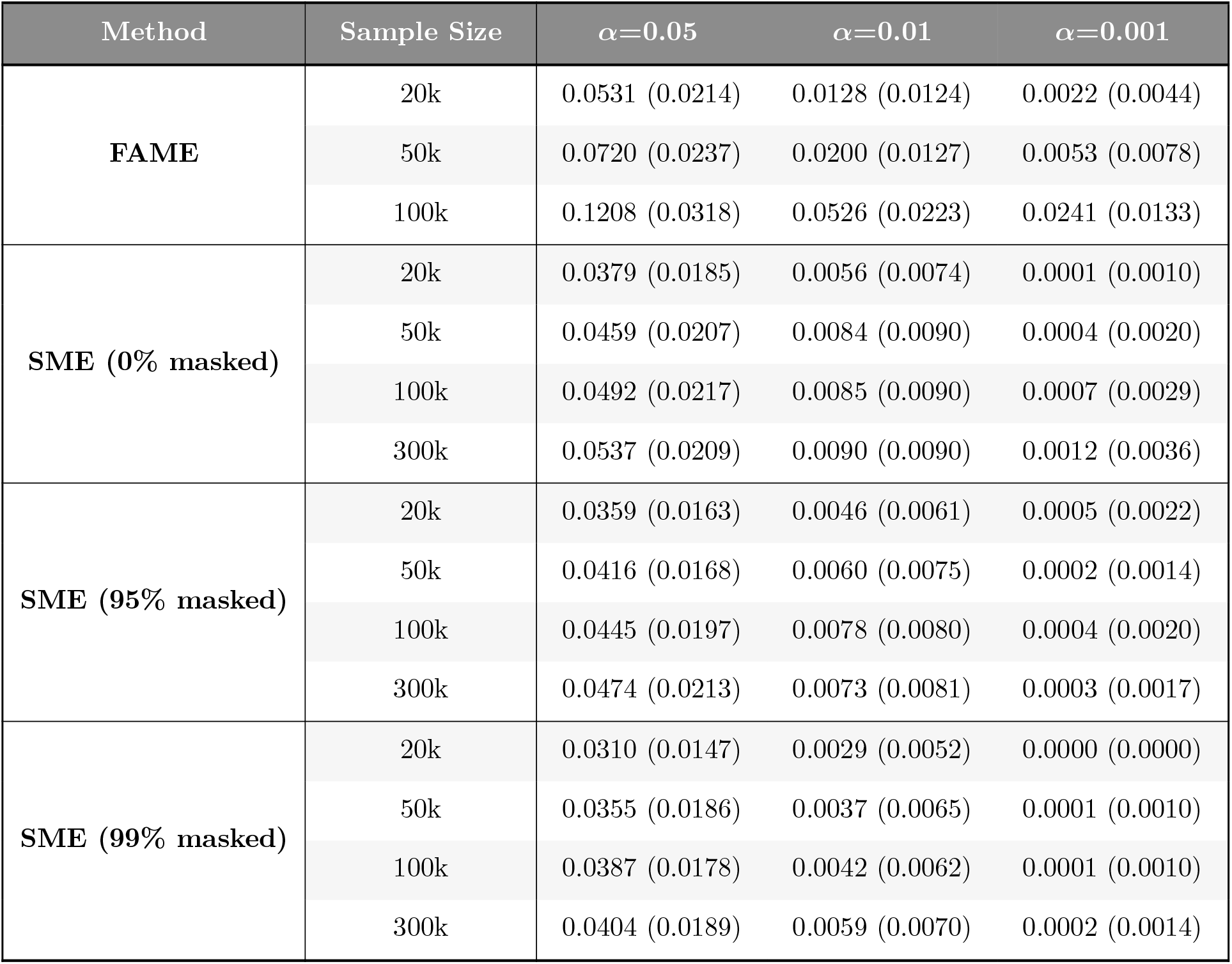
While using a mask that induces uniform sparsity, SME controls type I error rates when synthetic traits are generated under the null model. Synthetic traits were simulated with only additive effects using chromosome 1 from individuals of self-identified European ancestry in the UK Biobank. These data were then subsampled using sample sizes of 20k, 50k, 100k, and 300k individuals. A total of 100 causal additive variants were randomly selected for each trait and their effects were assumed to explain 40% of the phenotypic variance. Data were analyzed using both FAME (as a baseline) and SME under varying percentages of SNPs that are masked (0%, 95%, and 99%, respectively). Empirical size for the analyses used significance thresholds of *α* = 0.05, 0.01, and 0.001. Values in the parentheses are the standard deviations of the estimates. Results are based on 100 simulations per scenario. Due to computational constraints, the data with 300k individuals was only analyzed with SME.

**Figure 3.**
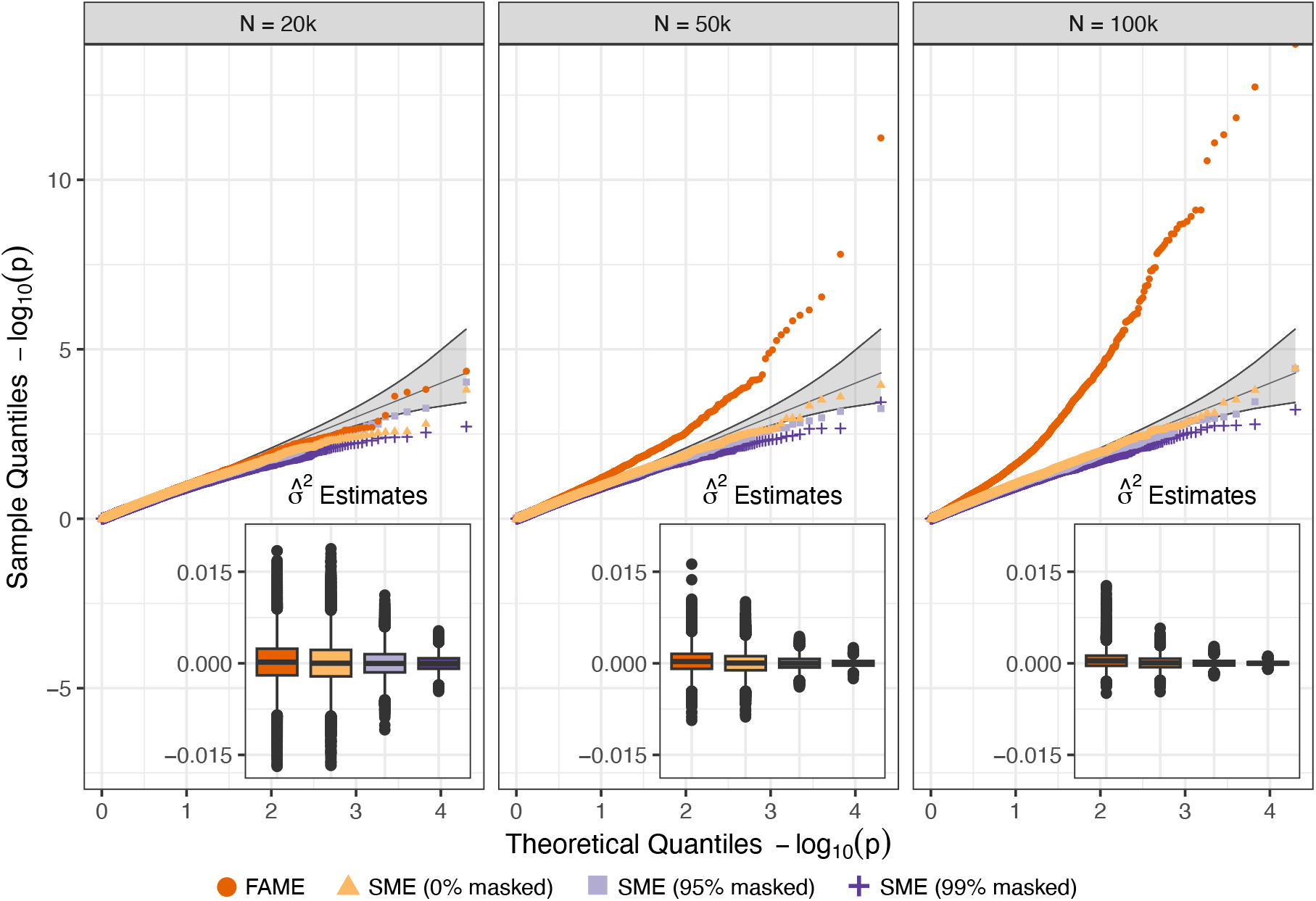
While using a mask that induces uniform sparsity, SME is well-calibrated under the null hypothesis and does not identify epistasis when traits are generated by only additive effects. Synthetic traits were simulated with only additive effects using chromosome 1 from individuals of self-identified European ancestry in the UK Biobank. These data were then subsampled using sample sizes of 20k, 50k, and 100k individuals. A total of 100 causal additive variants were randomly selected for each trait and their effects were assumed to explain 40% of the phenotypic variance. Data were analyzed using both FAME (as a baseline) and SME under varying percentages of SNPs that are masked (0%, 95%, and 99%, respectively). The small insets in each plot show the distribution of the estimated marginal epistatic variance components across all experiments. For reference, under the null hypothesis *H*_0_ : *σ*^2^ = 0. Results are based on 100 simulations per scenario.

Notably, using an external data source with a localized sparse masking scheme introduces a slight negative bias in variance component estimates produced by SME, leading to fewer significant *P*-values and more conservative inference (Fig. S5). While type I error control remains conservative (Table S1), this also means that the test may have reduced power when traits are indeed simulated under the alternative with non-zero epistatic effects. We will explore this behavior further in the next section.

### The masking strategy in SME leads to improved power in simulations

To assess the power of SME, we again generate synthetic continuous traits using real genotypes from chromosome 1 of white British individuals in the UK Biobank [2, 3, 5]. These data were subsampled using sample sizes of 50k, 100k, and 300k individuals. Here, we assume that 10% of all SNPs are causal and have additive effects that collectively explain 30% of the trait variance. Next, we fix the epistatic contribution to the trait variance to be 5%, making the total broad-sense heritability 35%. We select a set of epistatic variants from the causal SNPs and divide them into two equally sized groups. Each SNP in one group is simulated such that they only interact with SNPs in the other group. This simulation design gives control over the epistatic phenotypic variance explained (PVE) by the individual variants. In this analysis, we select 10, 20, 50, and 100 of the causal SNPs to be epistatic which corresponds to per SNP epistatic PVE equal to 1%, 0.5%, 0.2%, and 0.1% of the trait variance.

Once again, we analyze SME using two different external data source types that induce uniform and localized sparsity—masking out 0%, 95%, and 99% of the possible interactive partnerts when constructing the marginal epistatic covariance matrix **G**_*j*_ for each *j*-th focal SNP being tested (Fig. S4a). We compare the empirical power of SME to FAME as a baseline by assessing the respective abilities of both models to identify causal epsitatic SNPs at a genome-wide significance threshold *P* < 5 × 10^*-*8^ [36].

We find using the uniformly sparse masking scheme significantly enhances the power of SME, with greater levels of sparsity leading to better method performance (Fig. 4). When analyzing 300k individuals, the 99%-masked SME identifies at least 85.1% of the causal epistatic SNPs even when they contribute as little as 0.1% to the trait variance. This is compared to FAME and a non-masked SME which only detect at most 1% of causal SNPs with very small PVE. When epistatic variants have larger effect sizes and individually account for 1% of the trait variance, the 99%-masked SME shows 99.8% power even with relatively small sample sizes (e.g., 50k individuals). Again, this is compared to FAME and a non-masked SME which each only have approximately 35% power in this scenario. To examine the sensitivity of the SME to the specification of an external data source, we conducted a simulation in which the model was provided with a weight matrix that incorrectly masked true interacting partners [5]. Here, we observed that the SME framework protects against the false discovery of non-additive genetic effects and underestimates the marginal epistatic variance component (*σ*^2^) when causal SNPs involved in pairwise interactions were unobserved (Fig. S6).

**Figure 4.**
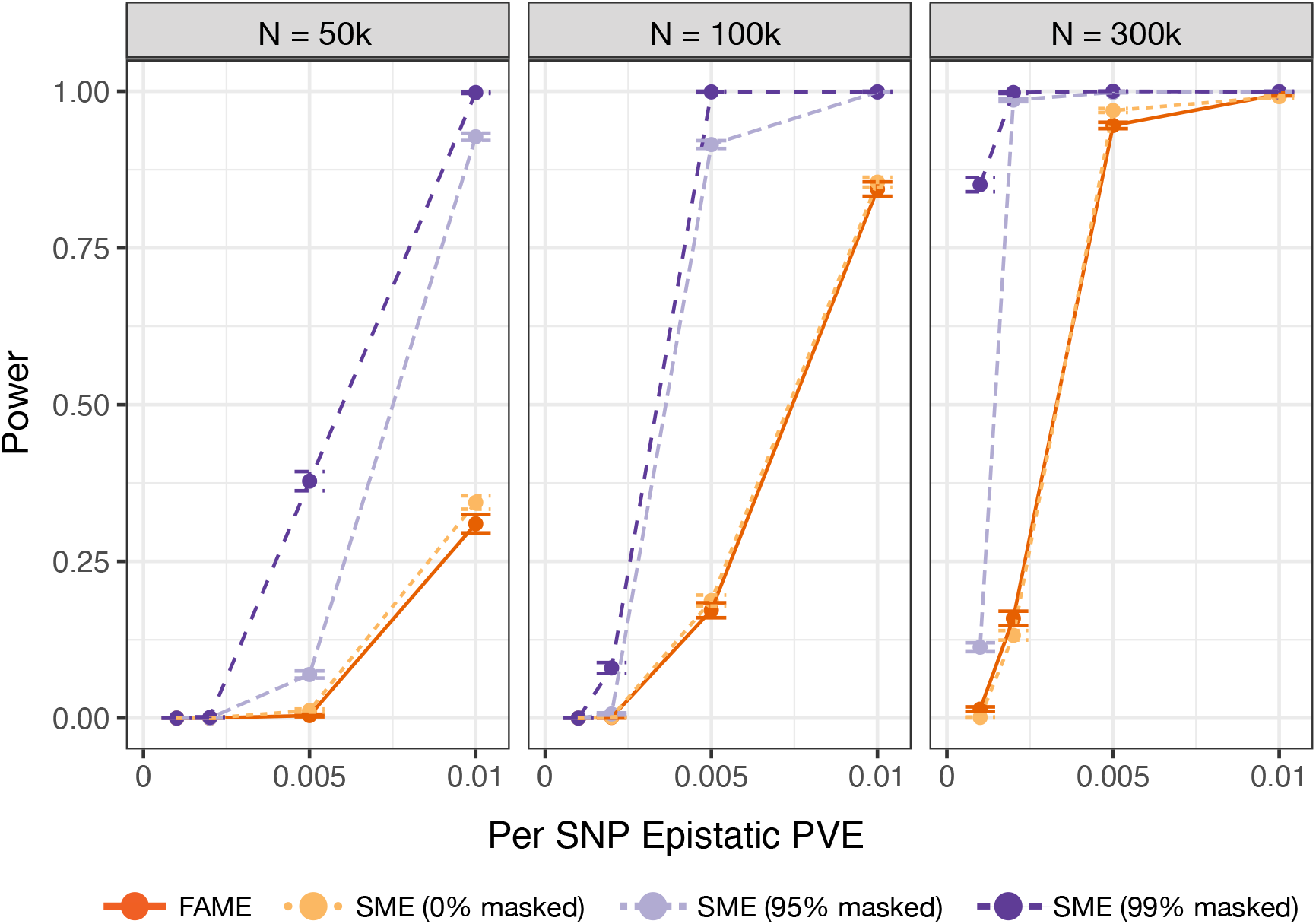
Uniform sparse modeling of interactions enhances the empirical power of SME. Synthetic traits were simulated with both additive and pairwise epistatic effects using chromosome 1 from individuals of self-identified European ancestry in the UK Biobank. Data were subsampled using sample sizes of 50k, 100k, and 300k individuals. We randomly selected 10% of all variants to have additive effects that collectively explained 30% of the trait variance. We then fixed the total epistatic variance to 5%. The per SNP epistatic phenotypic variance explained (PVE) was adjusted by varying the number of interacting SNPs (chosen to be 10, 20, 50, or 100 SNPs). Data were analyzed using both FAME (as a baseline) and SME under varying percentages of variants that are excluded from consideration as potential interaction partners for each focal SNP (0%, 95%, and 99% masking, respectively). Empirical power was determined using the significance threshold *P* < 5 ×10^*-*8^. Results are based on 100 simulations per scenario, with error bars representing the standard deviation across replicates.

Similar to the null simulation study, we see that SME produces negatively biased variance component estimates when using an external data source that induces localized sparsity in the model. Indeed, overconcentrating the search for potential interacting pairs to a select number of correlated variants leads to reduced empirical power compared to the masking resulting in uniform sparsity (Fig. S7). For example, when analyzing 300k individuals, the localized 99%-masked SME has just 6.5% power to identify epistatic variants that explain 0.1% of the trait variance. To overcome this issue in practice, we propose a strategy in which we take an external data source with localized genomic information and randomly unmask “unimportant” variants with uniform probability along the genome (essentially making the localized sparsity look more uniform as shown in Fig. S4b). As a demonstration of this idea, we implement a version of SME where we include an additional 1% and 5% of initially disregarded interactions back into the construction of **G**_*j*_ for each *j* = 1,. .., *J* tested focal SNPs in the data set. We find that adding this “noise” back into the mask reduces the negative bias of the variance component estimates and recovers as much as 28% of the power that was lost with respect to the uniformly sparse models (Fig. S8). For future users of the SME software, we want to note that there is a likely to be an application specific tradeoff between adding SNPs to a mask to reduce potential bias and finding the degree of sparsity needed for a optimally powered test.

### SME uses chromatin information to identify epistasis in hematology traits

We apply SME to four hematology traits — mean corpuscular hemoglobin (MCH), mean corpuscular hemoglobin concentration, mean corpuscular volume (MCV), and hematocrit (HCT) — assayed in 349,411 white British individuals in the UK Biobank [19] genotyped at 543,813 SNPs genome-wide. As an external data source, we leverage DNase I-hypersensitive sites (DHS) data measured over 12 days of *ex vivo* erythroid differentiation [34]. Of the quality controlled SNPs in our data, 4,932 of them are located in DHS regions enriched for transcriptional activity [28]. Since previous GWAS results have found genes associated with MCH, MCHC, and MCV to also be implicated in erythroid differentiation [37], we expect that conditioning SME to test over regulatory mechanisms gathered during erythropoiesis will be helpful in identifying epistatic variants for these traits. On the other hand, HCT is a phenotype that measures the percentage of red blood cells in an individual. Since the regulation for this trait has little to do with DHS sites, and more to do with oxygen available in the blood [38], we would expect a mask derived from functional data on erythropoiesis to not be helpful in enabling SME to detect epistasis.

For each trait, we use Manhattan plots to visually display the variant-level mapping results across each of the four traits, where chromosomes are shown in alternating colors for clarity (Figs. 5 and S9-S11). Corresponding genes that have SNPs with *P*-values below the genome-wide significance threshold to correct for multiple testing (*P* < 5 × 10^*-*8^) are also highlighted. Importantly, many of the marginal epistatic variants identified by SME are supported by multiple published studies that have investigated non-additive gene action related to erythropoiesis and red blood cell traits (Table 2).

**Table 2.** SME identifies marginal epistasis in hematology traits from individuals in the UK Biobank. Here, we analyze 349,411 white British individuals in the UK Biobank genotyped at 543,813 SNPs genome-wide. Traits in this analysis included: mean corpuscular hemoglobin (MCH), mean corpuscular hemoglobin concentration (MCHC), mean corpuscular volume (MCV), and hematocrit (HCT). As a mask, we leveraged DNase I-hypersensitive sites (DHS) data measured over 12 days of ex vivo erythroid differentiation [28, 34]. Listed are only results corresponding to SNPs that have marginal epistatic *P*-values below a genome-wide significance threshold to correct for multiple testing (*P* < 5 × 10^−8^). In the second and third columns, we list SNPs and their genomic location in the format `Chromosome:Basepair.` Next, we give the *P*-value and marginal epistatic phenotypic variance explained (PVE) for each SNP as esimtated by SME. The last columns detail the closest neighboring gene and a reference that have previously suggested some level of association or enrichment between each gene and the traits of interest.

| Trait | ID | Coordinates | $P$-Value | PVE | Gene | Ref. |
| --- | --- | --- | --- | --- | --- | --- |
| MCH | rs4711092 | 6:25684405 | $1.41 \times 10^{-11}$ | 0.007 | *SCGN* | [39, 45, 46] |
| MCH | rs9366624 | 6:25439492 | $1.80 \times 10^{-9}$ | 0.011 | *CARMIL1* | [37, 40, 45, 47, 48] |
| MCH | rs9461167 | 6:25418571 | $2.34 \times 10^{-9}$ | 0.007 | *CARMIL1* | [37, 40, 45, 47, 48] |
| MCH | rs9379764 | 6:25414023 | $5.53 \times 10^{-9}$ | 0.012 | *CARMIL1* | [37, 40, 45, 47–50] |
| MCH | rs441460 | 6:25548288 | $1.20 \times 10^{-8}$ | 0.008 | *CARMIL1* | [37, 40, 45, 47, 48] |
| MCH | rs198834 | 6:26114372 | $2.77 \times 10^{-8}$ | 0.008 | *H2BC4* | [51, 52] |
| MCH | rs13203202 | 6:25582771 | $3.17 \times 10^{-8}$ | 0.012 | *CARMIL1* | [37, 40, 45, 47, 48] |
| MCV | rs9276 | 6:33053577 | $9.09 \times 10^{-10}$ | 0.002 | *HLA-DPB1* | — |
| MCV | rs9366624 | 6:25439492 | $1.86 \times 10^{-8}$ | 0.008 | *CARMIL1* | [37, 40, 45, 47, 48] |

**Figure 5.**
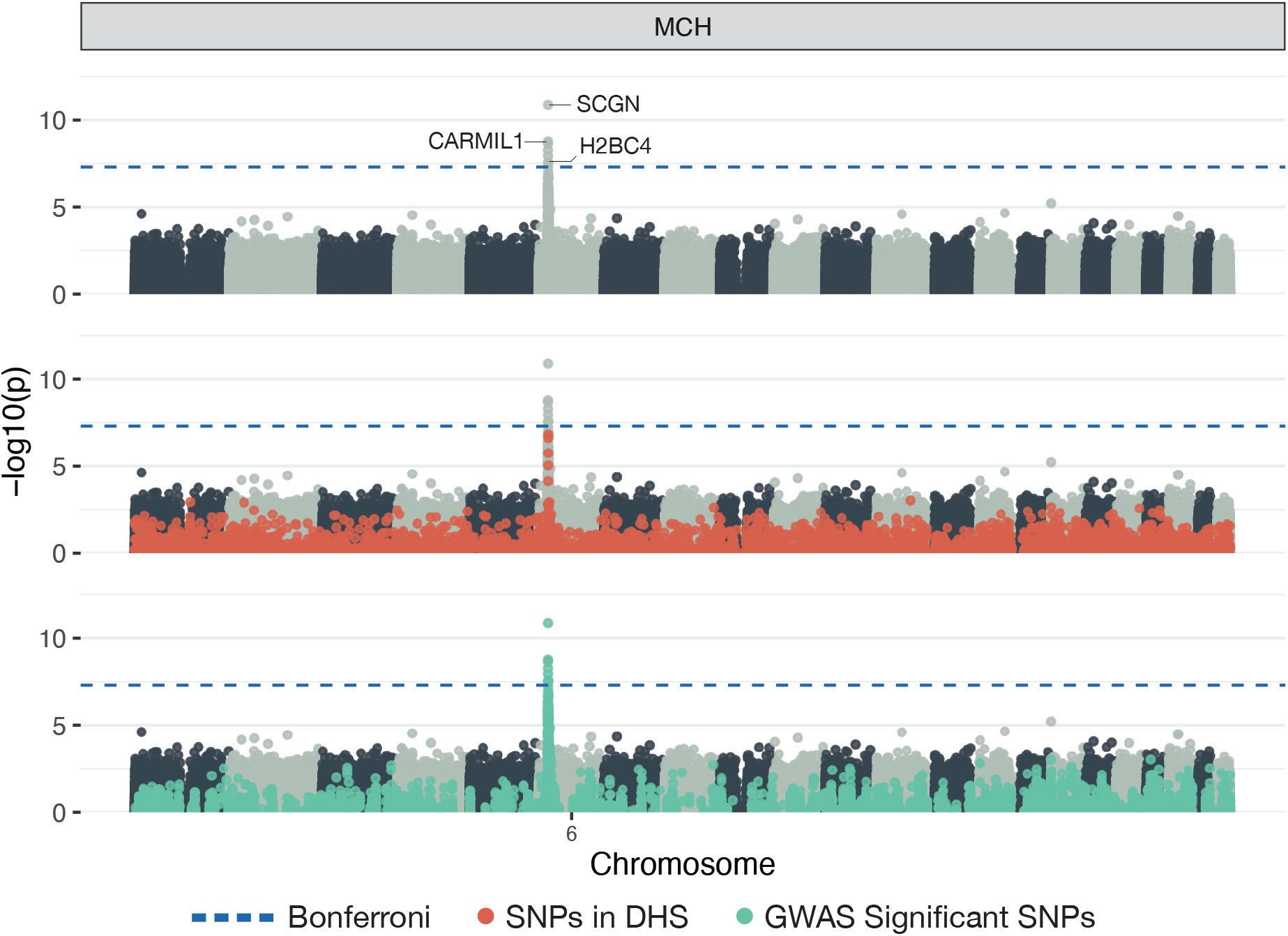
Manhattan plots of a genome-wide interaction analysis using SME to study mean corpuscular hemoglobin (MCH) assayed in individuals in the UK Biobank. As a mask in this study, we leveraged DNase I-hypersensitive sites (DHS) data measured over 12 days of *ex vivo* erythroid differentiation [28, 34]. This means that, while all SNPs are tested for marginal epistasis, only their interactions with SNPs in DHS regions are considered. Here, *-* log_10_ transformed *P*-values from SME are plotted for each SNP against their genomic positions. Chromosomes are shown in alternating colors for clarity. The dashed blue line represents the genome-wide significance threshold (*P* < 5 ×10^*-*8^). Each panel shows the same plot with different aspects of the result highlighted. The first simply shows the names of the closest neighboring genes to significant epistatic SNPs. The second panel highlights the SNPs that fall in DHS regions, and the third panel highlights SNPs that are also found to have a significant (additive) association with the trait according to a GWAS (Methods).

For example, when analyzing MCH, the strongest association identified by SME is the SNP rs4711092 (*P* = 1.41 × 10^*-*11^) maps to the gene secretagogin (*SCGN*) which regulates exocytosis by interacting with two soluble NSF adaptor proteins (*SNAP-25* and *SNAP-23*) and is critical for cell growth in some tissues [39]. For MCH, SME also identified five significantly associated SNPs (e.g., rs9366624 with *P* = 1.8 × 10^*-*9^) in the gene capping protein regulator and myosin 1 linker 1 (*CARMIL1*). *CARMIL1* is known to interact with and regulate the caping protein (*CP*) which plays a role via protein-protein interactions in regulating erythrpoiesis [40]. Specifically, *CARMIL* proteins regulate actin dynamics by regulating the activity of *CP* [41, 42]. Erythropoiesis leads to modifications in the expression of membrane and cytoskeletal proteins, whose interactions impact cell structure and function [43, 44]. Both genes *SCGN* and *CARMIL1* have previously been associated with hemoglobin concentration [37, 45]. A complete list of the results for all traits are listed in Tables S2-S5.

## Discussion

The marginal epistasis framework is an alternative to detect gene interactions. It derives its power by modeling the combined effect between a focal SNP and all other variants, thus alleviating the need to test every possible interaction separately. Still, current methods seeking to identify marginal epistasis struggle to scale to biobank-scale data and can be underpowered when non-additive genetic effects only explain a small portion of the overall trait variance [2, 3]. SME overcomes these limitations by inducing sparsity, essentially limiting the combined interaction for a focal SNP to just regions of the genome that have some known functional relationship with the trait of interest. This approach not only results in more efficient estimators but also offers a mechanism that allows the method to perform genome-wide analyses on modern data sets with runtimes that are magnitudes faster than previous approaches. Through extensive simulations, we show that SME controls type I error rates and produces calibrated *P*-values. We also show that SME has the power to detect SNPs involved in epistasis even when they explain very small fractions of the trait variance. By analyzing hematology traits from participants in the UK Biobank, we illustrate that SME, informed by DNase-seq data, identifies statistical epistasis in variants for which previous research have also found interaction pathways. We make SME available as open-source R software package to enable the broader community to easily use it in their research.

The current implementation of SME offers many directions for future development and applications. For example, the key to SME is that it relies on external data sources to induce sparsity in the model. While simulations show that misspecified or localized sparsity does not jeopardize the ability to control false positive rates, SME currently does not provide instructions on how to best format the external data for a particular analysis. In simulations, we show that some choices can induce a structure that leads to negative bias in the model estimates. We also show that adding random “noise” to the data can reduce this the bias. As part of future work, we will explore how to automatically balance this tradeoff within the software. Additionally, while SME uses an efficient model fitting algorithm, its current implementation has a non-negligible I/O overhead from repeatedly needing to read in (often large) genotype data into memory. Future development that optimizes this file read bottleneck has the potential to further improve the scalability of the method.

## Methods

### The sparse marginal epistasis test

Consider a genome-wide association study (GWAS) with *N* individuals who have been genotyped for *J* single nucleotide polymorphisms (SNPs) encoded as {0, 1, 2} copies of a reference allele at each locus. The marginal epistasis test aims to identify genetic variants that are involved in epistasis without exhaustively searching over all possible interactions [2]. SME refines this framework by incorporating functional genomic information into the model. Examining one SNP at a time (indexed by *J*), SME can be expressed via the following linear mixed model (LMM)

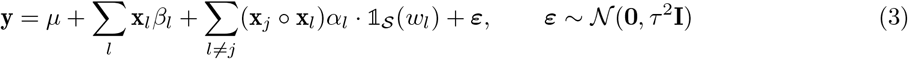

where **y** is an *N*-dimensional trait vector measured for each individual in the study; *µ* is an intercept term; **X** is the *N* × *J* matrix of genotypes with **x**_*l*_ representing an *N*-dimensional vector for the *l*-th SNP; *β*_*l*_ is the additive effect for the *l*-th SNP; **x**_*j*_ ○ **x**_*l*_ is the Hadamard (elementwise) product of the two genotypic vectors with corresponding interaction effect size *α*_*l*_; *ε* is a normally distributed error term with mean zero and scale variance term *τ*^2^; and **I** denotes an *N* × *N* identity matrix. The term 𝟙_𝒮_(*w*_*l*_) is an indicator function that encodes known regulatory information (or some other biological annotation) about the *l*-th SNP based on an external data source 𝒮. For example, using a DNase-seq experiment, *w*_*l*_ could denote whether the *l*-th SNP is located in a genomic region with chromatin accessibility (e.g., Fig. 1). Through this masking strategy, SME is able to leverage evidence from other studies to power its test for epistatic effects.

### Variance component model formulation

For biobank-scale data, there are often more SNPs than individuals. To overcome an undetermined system in Eq. (3), the marginal epistasis framework assumes that the effect sizes follow univariate normal distributions where *β*_*l*_ ∼ 𝒩 (0, *ω*^2^/*J*) and *α*_*l*_ ∼ 𝒩 (0, *σ*^2^/*J*^*^) with *J*\* =∑_*l*≠*j*_ 𝟙_𝒮_ (*w*_*l*_) representing the number of interactions considered in the model [2, 33, 53–55]. Assuming that the phenotype has been mean centered and scaled, these normal assumptions allow the LMM in Eq. (3) to be rewritten as

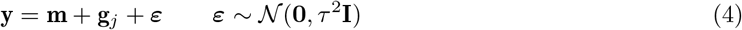

where **m** ∼ 𝒩 (**0**, *σ*^2^**K**) is the combined additive effects from all SNPs with relatedness matrix **K** = **XX**^**⊤**^/*J* and **g**_*j*_ ∼ 𝒩 (0, *σ*^2^**G**_*j*_) represents the effects of all pairwise interaction involving the *J*-th SNP that have not been masked. Here, we let the covariance 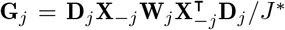 where **X**_*-j*_ denotes the genotype matrix without the *j*-th SNP and **D**_*j*_ = diag(**x**_*j*_) is an *N* × *N* diagonal matrix. Importantly, **W**_*j*_ = diag[𝟙_𝒮_(*w*_1_),. .., 𝟙_𝒮_(*w*_*J−*1_)] with binary diagonal elements that only equate to 1 if the *l*-th SNP satisfies the criteria from the external data source 𝒮.

### Point estimates

The model in Eq. (4) has three variance components that can be estimated using a computationally efficient method-of-moments (MoM) algorithm [23]. In expectation

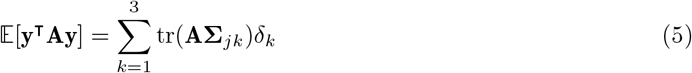

with **A** being a symmetric and non-negative definite matrix used to create weighted second moments, tr(•) denotes the trace of a matrix, and we use shorthand to represent [**∑**_*j*1_; **∑**_*j*2_; **∑**_*j*3_] = [**K**; **G**_*j*_; **I**] and ***δ***= (*ω*^2^, *σ*^2^, *τ*^2^), respectively. In practice, we replace the left hand side of Eq. (5) with the realized value **y**^**⊤**^**Ay**. We also use the realized covariance matrices in place of the arbitrary **A**. The point estimates for each variance component is then given as

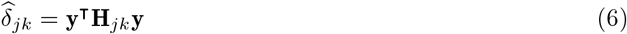

where 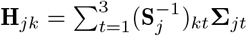 and **S**_*j*_ is a 3 × 3 matrix with elements (**S**_*j*_)_*kt*_ = tr(**∑**_*jk*_**∑**_*jt*_).

### Hypothesis testing

SME tests for non-zero marginal epistasis using a one-sides z-score or normal test. This is equivalent to assessing the null hypothesis *H*_0_ : *σ*^2^ ≤ 0 for each SNP in the data. We derive a test statistic with the estimate 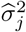 using Eq. (6) and compute an approximate standard error

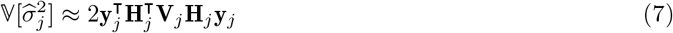

where 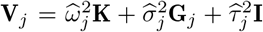. Note that the point estimates from Eqs. (5) and (6) are unbiased and can lead to negative values when the true variance component is zero [23]. The one-sided hypothesis test formalizes the constraint that only positive estimates of 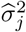 can be indicative of marginal epistasis.

### Scalable computation via stochastic MoM

The right-hand side of Eq. (6) involves computing traces of matrix products. If each covariance matrix is held in memory, then this can be done efficiently (without matrix multiplication) using the Frobenius inner product. However, holding large covariance matrices in memory itself prevents scalability of the method. Naively multiplying two *N* × *N* matrices requires *N*^3^ field operations. This too can be impractical for biobank-scale data with large sample sizes. To enable genome-wide testing, SME makes use of a stochastic MoM approach through the implementation of Hutchinson’s stochastic trace estimator and the Mailman algorithm.

### Hutchinson’s stochastic trace estimator

Hutchinson’s stochastic trace estimator approximates the trace of a matrix product via the following [4, 24, 26, 35]

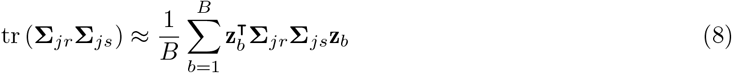

where **z**_*b*_ ∼ 𝒩 (**0, I**) is a normally distributed vector and *B* is the number of random draws used to approximate the trace. This operation only depends on a series of matrix-by-vector products and has time complexity 𝒪 (*BNJ*). Essentially, we choose an order of operations such that computing the quadratic forms 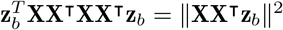 is reduced to (i) applying **X**^**⊤**^ to the vector **z**_*b*_ and then (ii) applying X to the resulting vector 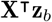 for all *B* random vectors. Importantly, Eq. (8) can be set up algorithmically such that the approximation is done block-wise over the individual-level genotype data (Supplementary Material). Using this approach, SME avoids having to compute any of the covariance matrices directly and alleviates the need to load the entire genotype matrix into memory all at once.

### The Mailman algorithm

An additional computational speedup can be achieved by making use of the discrete encoding for each SNP. The Mailman algorithm allows for an *N* × *J* matrix to be multiplied by any real vector in 𝒪 (*NJ*/log_Ω_ (max{*N, J*})) time if it has elements defined over a finite alphabet size Ω [4, 24, 27, 35]. A standardized genotype matrix can be written as **X** = (**A − U**)**Q**^*-*1^ where **A** is an *N* × *J* allele count matrix with elements *a*_*ij*_ ∈ {0, 1, 2} over finite size Ω = 3, **U** = [**u**_1_,…, **u**_*J*_] is a matrix where the *j*-th column contains the sample mean for the *j*-th SNP, and 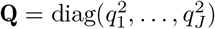 contains the variance of each SNP as the diagonal entries. With this specification, we can write 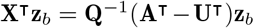.The first term 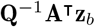 can be solved in 𝒪 (*NJ*/log_3_(max{*N, J*})) time. The second term 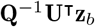 corresponds to scaling the random *N*-dimensional vector **z**_*b*_ which can be computed in time 𝒪 (*N* + *J*) [4, 24, 35].

### Shared random vectors and parallelization

With the stochastic trace estimator and the Mailman algorithm, it is feasible to estimate the variance components for each focal SNP even when a study has a large number of individuals. Still, testing every focal SNP against all variants genome-wide remains a challenge. In SME, we propose randomly selecting subsets of focal SNPs and having them share the same random vectors **z**_*b*_ when performing the stochastic trace estimation. This limits the number of computations that need to be performed while maintaining unbiasedness in the point estimates (Fig. S12).

Since the error terms in Eq. (4) are assumed to be independent, the only two intermediate products that need to be computed in Eq. (8) are 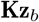 and 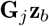.With these terms, we can compute all combinations of traces of matrix products that are required to fit SME (e.g.,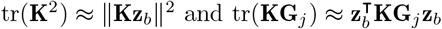). For a subset of *L* focal SNPs and fixed **z**_*b*_, the term 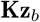 is constant and only 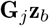 changes. This reduces the effective time needed to compute 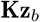 per test by a factor of 1/*L*. Fig. S1 illustrates the idea of sharing random vectors.

As previously mentioned, reading biobank-scale genotype data into memory requires non-negligible over-head. The `R` implementation of SME reads in genotypes once for each subset of focal SNPs that share the same random vectors. The computation of 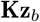 and each 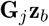 can then be done in parallel using multithreading.

### Masking further reduces the effective size of data

A direct benefit of masking is that it is equivalent to removing entire columns from the genotype matrix. This reduction contributes to a significantly faster runtime when computing 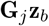 (e.g., Figs. S2 and S3). Concretely, applying a mask to an *N* × *J* genotype matrix reduces the number of columns to *J*\* ≤ *J*. If the mask is sparse enough such that *J*\* ≤ *N*, the time complexity of the Mailman algorithm is also reduced to 𝒪 (*BNJ*\* /log_3_(*N*)).

### Preprocessing the UK Biobank

Genotype data for 488,377 individuals in the UK Biobank were downloaded and converted using the `ukbgene` and `ukbconv` tools, respectively. Continuous traits were also downloaded using the ukbgene tool and were adjusted for age and sex. Individuals identified as having high heterozygosity, excessive relatedness, or aneuploidy were removed (1,550 individuals). After separating individuals into self-identified ancestral cohorts using data field `21000`, unrelated individuals were selected by randomly choosing one person from each related pair. This resulted in *N* = 349,411 white British individuals to be included in our analysis. We downloaded imputed SNP data from the UK Biobank for all remaining individuals and removed SNPs with an information score below 0.8. Information scores for each SNP are provided by the UK Biobank (http://biobank.ctsu.ox.ac.uk/crystal/refer.cgi?id=1967).

Quality control for the remaining genotyped and imputed 1,933,118 variants was then performed on each cohort separately using the following steps. All structural variants were first removed, leaving only single nucleotide polymorphisms (SNPs) in the genotype data. Next, all AT/CG SNPs were removed to avoid possible confounding due to sequencing errors. Then, SNPs with minor allele frequency less than 1% were removed using the `PLINK 2.0` [56] command `--maf 0.01`. We then removed all SNPs found to be out of Hardy-Weinberg equilibrium, using the `PLINK --hwe 0.000001` flag to remove all SNPs with a Fisher’s exact test *P*-value < 10^*-*6^. Finally, SNPs with any missingness were removed using the `PLINK 2.0 --geno 0.00` flag. This left a total of *J* = 543,813 SNPs for our study.

### Generating masks from external data sources

#### Masks using DNase I hypersensitive sites

The chromatin accessibility-based masks were derived from DNase I hypersensitive sites (DHS) measured over 12 days of *ex vivo* erythroid differentiation [34]. The DHS intervals were reported using the hg38 human reference genome. To map correspondence to the UK Biobank data (which uses the hg19 as reference), we performed a lift-over using `CrossMap` [57]. We mapped each SNP in the UK Biobank to the genomic intervals in the DHS data using the R software package `GenomicRanges` [58]. The resulting mask comprised of *J*\* = 4952 SNPs.

#### Small note on linkage disequilibrium blocks

To control the type I error rate, variants in the same linkage disequilibrium (LD) block as the *j*-th focal SNP are also masked. The LD blocks used for this study were approximately independent and derived using European individuals [59].

### GWAS summary statistics

Summary statistics used to compare against marginal epistatic results for each trait in the UK Biobank were downloaded from https://www.nealelab.is/uk-biobank. These summary statistics were first filtered to match the same set of SNPs that passed our quality control. SNPs that were reported as being associated with a trait at genome-wide significance (*P* < 5 × 10^*-*8^) in the UK Biobank European cohort were highlighted in Figs. 5 and S9-S11.

### Simulation studies

To characterize the behavior of the sparse marginal epistasis test, we generate quantitative traits using chromosome 1 of the white British cohort from the UK Biobank [2, 3, 5]. This data consisted of *N* = 349,411 individuals and *J* = 43,332 SNPs. Here, we sample 10% of the SNPs in the data to be causal and simulate traits using the following linear model

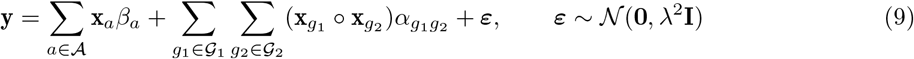

where **y** is an *N*-dimensional phenotype vector; **𝒜** represents the set of causal SNPs with additive effects; **x**_*a*_ is the genotype for the *a*-th causal SNP encoded as 0, 1, or 2 copies of a reference allele;_*a*_ is the additive effect sizes for the *a*-th SNP; both 𝒢_1_ and 𝒢_2_ are sets of epistatic SNPs that are non-overlapping subsets of 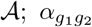 is the interaction effect sizes between 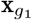 and 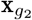; and ε is a vector of normally distributed environmental noise. We sample the effect sizes from standard normal distributions and rescale them so that the additive and epistatic effects explain a desired proportion of the trait variance. Specifically, the additive and epistatic variance components make up the broad-sense heritability of the trait 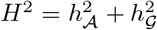. Similarly, the environmental noise matrix is also rescaled such that it explains the remaining 1 *-H*^2^ proportion of the trait variance.

In this simulation design, the epistatic causal SNPs interact between sets. All SNPs in 𝒢_1_ interact with all SNPs in the 𝒢_2_, but do not interact with variants in their own group (and vice versa). Note that we use this setup because of the ability to detect interacting variants in the marginal epistasis framework depends on the proportion of phenotypic variance that they marginally explain. The parameters that let us control the phenotypic variance explained by a single SNP are the epistatic heritability 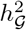 and the cardinality of the set that a SNP belongs to. For example, a SNP in 𝒢_1_ will explain on average 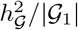 of the total phenotypic variance.

To simulate masks, we select some proportion of the non-epistatic SNPs to zero out of the interaction covariance matrix. For example, when analyzing the *j*-th SNP, 95% masking corresponds to excluding 41,165 out of the 43,332 SNPs when computing **G**_*j*_. Similarly, only 433 SNPs are used when using a 99% masking strategy. To create masks that induce “uniform sparsity”, we randomly sample from all SNPs in the data set with uniform probability. In real data, masks are likely not sampled uniformly. An obvious potential source of complication for non-uniform masking can come from linkage disequilibrium between SNPs. Therefore, to simulate masks which induce “localized sparsity”, we randomly sample a seed SNP and define a genomic window around it. SNPs outside that window are then masked.

## Data and software availability

Source code and tutorials for implementing the “sparse marginal epistasis test” (SME) are publicly available as an R package which is available online at https://github.com/lcrawlab/sme. Software for running the “FAst Marginal Epistasis test” (FAME) is freely available at https://github.com/sriramlab/FAME. The original “MArginal ePIstasis Test” (MAPIT) was implemented using the `mvmapit` software package in R and is available both on CRAN (https://cran.r-project.org/package=mvMAPIT) and GitHub (https://github.com/lcrawlab/mvMAPIT). All software for SME, FAME, and MAPIT were fit using their default settings, unless otherwise stated in the main text. Data from the UK Biobank Resource was made available under Application Numbers 14649 and can be accessed by direct application to the UK Biobank. GWAS summary statistics were downloaded from https://www.nealelab.is/uk-biobank and corresponding linkage disequilibrium maps were taken from https://bitbucket.org/nygcresearch/ldetect-data/. The DNase I hypersensitive site (DHS) data are available at https://doi.org/10.5281/zenodo.5291736. We used CrossMap (https://crossmap.readthedocs.io/) and `GenomicRanges` (https://doi.org/doi:10.18129/B9.bioc.GenomicRanges) to map SNPs from the UK Biobank to genomic intervals in the DHS data. The chain file for the `CrossMap` liftover tool can be found at http://hgdownload.soe.ucsc.edu/goldenPath/hg38/liftOver/. The full list of summary statistics from the genome-wide interaction analysis using SME to study hematology traits in UK Biobank are publicly available at https://doi.org/10.5281/zenodo.14607998.

## Acknowledgments

We thank members of the Weinreich and Crawford Labs for insightful comments on earlier versions of this manuscript as well as Bogdan Pasaniuc (University of Pennsylvania) and Roberta DeVito (Brown University) for helpful discussions. We also thank Ashok Ragavendran and the Computational Biology Core (CBC) for advice on software development. This research was conducted in part using computational resources and services at the Center for Computation and Visualization at Brown University. This research was also conducted using the UK Biobank Resource under Application Numbers 14649.

## Author contributions

JS and LC conceived the study. JS and LC developed the methods. SPS preprocessed and provided the data. JS developed the software and performed the analyses. DW and LC supervised the project and provided resources. All authors wrote and revised the manuscript.

## References

1. Abdellaoui, A., Yengo, L., Verweij, K. J. H. & Visscher, P. M. 15 years of GWAS discovery: Realizing the promise. The American Journal of Human Genetics 110, 179–194. issn: 0002-9297. https://www.sciencedirect.com/science/article/pii/S0002929722005456 (2024) (Feb. 2023).

2. Crawford, L., Zeng, P., Mukherjee, S. & Zhou, X. Detecting epistasis with the marginal epistasis test in genetic mapping studies of quantitative traits. en. PLOS Genetics 13 (ed Lin, X.) e1006869. issn: 1553-7404. 10.1371/journal.pgen.1006869 (2021) (July 2017).

3. Stamp, J., DenAdel, A., Weinreich, D. & Crawford, L. Leveraging the Genetic Correlation between Traits Improves the Detection of Epistasis in Genome-wide Association Studies. G3 Genes|Genomes|Genetics, jkad118. issn: 2160-1836. 10.1093/g3journal/jkad118 (2023) (May 2023).

4. Fu, B. et al. A biobank-scale test of marginal epistasis reveals genome-wide signals of polygenic epistasis en. Sept. 2023. https://www.biorxiv.org/content/10.1101/2023.09.10.557084v1 (2024).

5. Pattillo Smith, S. et al. Discovering non-additive heritability using additive GWAS summary statistics. eLife 13 (ed Perry, G. H.) e90459. issn: 2050-084X. 10.7554/eLife.90459 (2024) (June 2024).

6. Balvert, M. et al. Considerations in the search for epistasis. Genome Biology 25, 296. issn: 1474-760X. 10.1186/s13059-024-03427-z (2024) (Nov. 2024).

7. Weinreich, D. M., Delaney, N. F., DePristo, M. A. & Hartl, D. L. Darwinian Evolution Can Follow Only Very Few Mutational Paths to Fitter Proteins. Science 312, 111–114. https://www.science.org/doi/10.1126/science.1123539 (2021) (Apr. 2006).

8. Fröhlich, C. et al. Epistasis arises from shifting the rate-limiting step during enzyme evolution of a - lactamase. en. Nature Catalysis 7, 499–509. issn: 2520-1158. https://www.nature.com/articles/s41929-024-01117-4 (2024) (May 2024).

9. Mackay, T. F. C. & Anholt, R. R. H. Pleiotropy, epistasis and the genetic architecture of quantitative traits. en. Nature Reviews Genetics, 1–19. issn: 1471-0064. https://www.nature.com/articles/s41576-024-00711-3 (2024) (Apr. 2024).

10. Weatherly, S. M. et al. Identification of Arhgef12 and Prkci as genetic modifiers of retinal dysplasia in the Crb1rd8 mouse model. en. PLOS Genetics 18, e1009798. issn: 1553-7404. https://journals.plos.org/plosgenetics/article?id=10.1371/journal.pgen.1009798 (2024) (June 2022).

11. Zwarts, L. et al. Complex genetic architecture of Drosophila aggressive behavior. Proceedings of the National Academy of Sciences 108, 17070–17075. https://www.pnas.org/doi/full/10.1073/pnas.1113877108 (2024) (Oct. 2011).

12. Polderman, T. J. C. et al. Meta-analysis of the heritability of human traits based on fifty years of twin studies. en. Nature Genetics 47, 702–709. issn: 1546-1718. https://www.nature.com/articles/ng.3285 (2024) (July 2015).

13. Hemani, G. et al. Phantom epistasis between unlinked loci. en. Nature 596, E1–E3. issn: 1476-4687. https://www.nature.com/articles/s41586-021-03765-z (2024) (Aug. 2021).

14. Hivert, V. et al. Estimation of non-additive genetic variance in human complex traits from a large sample of unrelated individuals. en. The American Journal of Human Genetics 108, 786–798. issn: 0002-9297. https://www.sciencedirect.com/science/article/pii/S0002929721000562 (2022) (May 2021).

15. Purcell, S. et al. PLINK: A Tool Set for Whole-Genome Association and Population-Based Linkage Analyses. American Journal of Human Genetics 81, 559–575. issn: 0002-9297. https://www.ncbi.nlm.nih.gov/pmc/articles/PMC1950838/ (2021) (Sept. 2007).

16. Schüpbach, T., Xenarios, I., Bergmann, S. & Kapur, K. FastEpistasis: a high performance computing solution for quantitative trait epistasis. Bioinformatics 26, 1468–1469. issn: 1367-4803. https://www.ncbi.nlm.nih.gov/pmc/articles/PMC2872003/ (2021) (June 2010).

17. Prabhu, S. & Pe’er, I. Ultrafast genome-wide scan for SNP–SNP interactions in common complex disease. en. Genome Research 22, 2230–2240. issn: 1088-9051, 1549-5469. https://genome.cshlp.org/content/22/11/2230 (2021) (Nov. 2012).

18. Wan, X. et al. BOOST: A Fast Approach to Detecting Gene-Gene Interactions in Genome-wide Case-Control Studies. American Journal of Human Genetics 87, 325–340. issn: 0002-9297. https://www.ncbi.nlm.nih.gov/pmc/articles/PMC2933337/ (2021) (Sept. 2010).

19. Bycroft, C. et al. The UK Biobank resource with deep phenotyping and genomic data. en. Nature 562, 203–209. issn: 1476-4687. https://www.nature.com/articles/s41586-018-0579-z (2022) (Oct. 2018).

20. Nagai, A. et al. Overview of the BioBank Japan Project: Study design and profile. Journal of Epidemiology 27, S2–S8. issn: 0917-5040. https://www.ncbi.nlm.nih.gov/pmc/articles/PMC5350590/ (2024) (Feb. 2017).

21. Moore, R. et al. A linear mixed-model approach to study multivariate gene–environment interactions. en. Nature Genetics 51, 180–186. issn: 1546-1718. https://www.nature.com/articles/s41588-018-0271-0 (2025) (Jan. 2019).

22. Haseman, J. K. & Elston, R. C. The investigation of linkage between a quantitative trait and a marker locus. en. Behavior Genetics 2, 3–19. issn: 1573-3297. 10.1007/BF01066731 (2024) (Mar. 1972).

23. Zhou, X. A unified framework for variance component estimation with summary statistics in genome-wide association studies. EN. Annals of Applied Statistics 11, 2027–2051. issn: 1932-6157, 1941-7330. https://projecteuclid.org/euclid.aoas/1514430276 (2021) (Dec. 2017).

24. Wu, Y. & Sankararaman, S. A scalable estimator of SNP heritability for biobank-scale data. Bioinformatics 34, i187–i194. issn: 1367-4803. 10.1093/bioinformatics/bty253 (2021) (July 2018).

25. Mbatchou, J. et al. Computationally efficient whole-genome regression for quantitative and binary traits. en. Nature Genetics 53, 1097–1103. issn: 1546-1718. https://www.nature.com/articles/s41588-021-00870-7 (2025) (July 2021).

26. Hutchinson, M. A Stochastic Estimator of the Trace of the Influence Matrix for Laplacian Smoothing Splines. en. Communications in Statistics - Simulation and Computation 18, 1059–1076. issn: 0361-0918, 1532-4141. http://www.tandfonline.com/doi/abs/10.1080/03610918908812806 (2024) (Jan. 1989).

27. Liberty, E. & Zucker, S. W. The Mailman algorithm: a note on matrix vector multiplication. en.

28. Maurano, M. T. et al. Systematic Localization of Common Disease-Associated Variation in Regulatory DNA. Science 337, 1190–1195. https://www.science.org/doi/10.1126/science.1222794 (2022) (Sept. 2012).

29. Finucane, H. K. et al. Partitioning heritability by functional annotation using genome-wide association summary statistics. en. Nature Genetics 47, 1228–1235. issn: 1546-1718. https://www.nature.com/articles/ng.3404 (2023) (Nov. 2015).

30. Cano-Gamez, E. & Trynka, G. From GWAS to Function: Using Functional Genomics to Identify the Mechanisms Underlying Complex Diseases. English. Frontiers in Genetics 11. issn: 1664-8021. https://www.frontiersin.org/articles/10.3389/fgene.2020.00424/full (2021) (2020).

31. Finucane, H. K. et al. Heritability enrichment of specifically expressed genes identifies diseaserelevant tissues and cell types. en. Nature Genetics 50, 621–629. issn: 1546-1718. https://www.nature.com/articles/s41588-018-0081-4 (2024) (Apr. 2018).

32. Boix, C. A., James, B. T., Park, Y. P., Meuleman, W. & Kellis, M. Regulatory genomic circuitry of human disease loci by integrative epigenomics. en. Nature 590, 300–307. issn: 1476-4687. https://www.nature.com/articles/s41586-020-03145-z (2022) (Feb. 2021).

33. Yang, J. et al. Common SNPs explain a large proportion of the heritability for human height. en. Nature Genetics 42, 565–569. issn: 1546-1718. https://www.nature.com/articles/ng.608 (2021) (July 2010).

34. Georgolopoulos, G. et al. Discrete regulatory modules instruct hematopoietic lineage commitment and differentiation. en. Nature Communications 12, 6790. issn: 2041-1723. https://www.nature.com/articles/s41467-021-27159-x (2024) (Nov. 2021).

35. Wu, Y. et al. Fast estimation of genetic correlation for biobank-scale data. American Journal of Human Genetics 109, 24–32. issn: 0002-9297. https://www.ncbi.nlm.nih.gov/pmc/articles/PMC8764132/ (2024) (Jan. 2022).

36. Fadista, J., Manning, A. K., Florez, J. C. & Groop, L. The (in)famous GWAS P-value threshold revisited and updated for low-frequency variants. en. European Journal of Human Genetics 24, 1202–1205. issn: 1476-5438. https://www.nature.com/articles/ejhg2015269 (2025) (Aug. 2016).

37. Ding, K. et al. Genetic Loci Implicated in Erythroid Differentiation and Cell Cycle Regulation Are Associated With Red Blood Cell Traits. Mayo Clinic Proceedings 87, 461–474. issn: 0025-6196. https://www.ncbi.nlm.nih.gov/pmc/articles/PMC3538470/ (2024) (May 2012).

38. Bhoopalan, S. V., Huang, L. J.-s. & Weiss, M. J. Erythropoietin regulation of red blood cell production: from bench to bedside and back. en. F1000Research 9, F1000 Faculty Rev. https://pmc.ncbi.nlm.nih.gov/articles/PMC7503180/ (2024) (Sept. 2020).

39. Qin, J. et al. Structural and mechanistic insights into secretagogin-mediated exocytosis. Proceedings of the National Academy of Sciences 117, 6559–6570. https://www.pnas.org/doi/10.1073/pnas.1919698117 (2024) (Mar. 2020).

40. Ray, S. et al. Functional requirements for a Samd14-capping protein complex in stress erythropoiesis. eLife 11, e76497. issn: 2050-084X. 10.7554/eLife.76497 (2024) (June 2022).

41. Jung, G., Pan, M., Alexander, C. J., Jin, T. & Hammer, J. A. Dual regulation of the actin cytoskeleton by CARMIL-GAP. Journal of Cell Science 135, jcs258704. issn: 0021-9533. 10.1242/jcs.258704 (2024) (June 2022).

42. Stark, B. C., Lanier, M. H. & Cooper, J. A. CARMIL family proteins as multidomain regulators of actin-based motility. Molecular Biology of the Cell 28, 1713–1723. issn: 1059-1524. https://www.ncbi.nlm.nih.gov/pmc/articles/PMC5491179/ (2024) (July 2017).

43. Nigra, A. D., Casale, C. H. & Santander, V. S. Human erythrocytes: cytoskeleton and its origin. en. Cellular and Molecular Life Sciences 77, 1681–1694. issn: 1420-9071. 10.1007/s00018-019-03346-4 (2024) (May 2020).

44. Gokhin, D. S. & Fowler, V. M. Feisty filaments: actin dynamics in the red blood cell membrane skeleton. en-US. Current Opinion in Hematology 23, 206. issn: 1065-6251. https://journals.lww.com/co-hematology/fulltext/2016/05000/feisty_filamentsactin_dynamics_in_the_red_blood.5.aspx (2024) (May 2016).

45. Timoteo, V. J., Chiang, K.-M.Yang, H.-C. & Pan, W.-H. Common and ethnic-specific genetic determinants of hemoglobin concentration between Taiwanese Han Chinese and European Whites: findings from comparative two-stage genome-wide association studies. The Journal of Nutritional Biochemistry 111, 109126. issn: 0955-2863. https://www.sciencedirect.com/science/article/pii/S0955286322001942 (2024) (Jan. 2023).

46. Bauer, M. C. et al. Identification of a high-affinity network of secretagogin-binding proteins involved in vesicle secretion. en. Molecular BioSystems 7, 2196–2204. issn: 1742-2051. https://pubs.rsc.org/en/content/articlelanding/2011/mb/c0mb00349b (2024) (Jan. 2011).

47. Edwards, M., Liang, Y., Kim, T. & Cooper, J. A. Physiological role of the interaction between CARMIL1 and capping protein. Molecular Biology of the Cell 24, 3047–3055. issn: 1059-1524. https://www.molbiolcell.org/doi/10.1091/mbc.e13-05-0270 (2024) (Oct. 2013).

48. Yang, C. et al. Mammalian CARMIL Inhibits Actin Filament Capping by Capping Protein. Developmental Cell 9, 209–221. issn: 1534-5807. https://www.sciencedirect.com/science/article/pii/S1534580705002510 (2024) (Aug. 2005).

49. Plotnikov, D. et al. High Blood Pressure and Intraocular Pressure: A Mendelian Randomization Study. eng. Investigative Ophthalmology & Visual Science 63, 29. issn: 1552-5783 (June 2022).

50. Galesloot, T. E. et al. Meta-GWAS and Meta-Analysis of Exome Array Studies Do Not Reveal Genetic Determinants of Serum Hepcidin. en. PLOS ONE 11, e0166628. issn: 1932-6203. https://journals.plos.org/plosone/article?id=10.1371/journal.pone.0166628 (2024) (Nov. 2016).

51. Vuckovic, D. et al. The Polygenic and Monogenic Basis of Blood Traits and Diseases. Cell 182, 1214–1231.e11. issn: 0092-8674. https://www.sciencedirect.com/science/article/pii/S0092867420309995 (2024) (Sept. 2020).

52. Zhang, W. et al. Evaluation of Genetic Variation Contributing to Differences in Gene Expression between Populations. English. The American Journal of Human Genetics 82, 631–640. issn: 0002-9297, 1537-6605. https://www.cell.com/ajhg/abstract/S0002-9297(08)00136-5 (2024) (Mar. 2008).

53. Bulik-Sullivan, B. K. et al. LD Score regression distinguishes confounding from polygenicity in genome-wide association studies. en. Nature Genetics 47, 291–295. issn: 1546-1718. https://www.nature.com/articles/ng.3211 (2022) (Mar. 2015).

54. Wu, M. C. et al. Rare-Variant Association Testing for Sequencing Data with the Sequence Kernel Association Test. American Journal of Human Genetics 89, 82–93. issn: 0002-9297. https://www.ncbi.nlm.nih.gov/pmc/articles/PMC3135811/ (2021) (July 2011).

55. Zhou, X., Carbonetto, P. & Stephens, M. Polygenic Modeling with Bayesian Sparse Linear Mixed Models. en. PLOS Genetics 9, e1003264. issn: 1553-7404. https://journals.plos.org/plosgenetics/article?id=10.1371/journal.pgen.1003264 (2021) (Feb. 2013).

56. Chang, C. C. et al. Second-generation PLINK: rising to the challenge of larger and richer datasets. eng. GigaScience 4, 7. issn: 2047-217X (2015).

57. Zhao, H. et al. CrossMap: a versatile tool for coordinate conversion between genome assemblies. Bioinformatics 30, 1006–1007. issn: 1367-4803. 10.1093/bioinformatics/btt730 (2024) (Apr. 2014).

58. Lawrence, M. et al. Software for Computing and Annotating Genomic Ranges. en. PLOS Computational Biology 9, e1003118. issn: 1553-7358. https://journals.plos.org/ploscompbiol/article?id=10.1371/journal.pcbi.1003118 (2024) (Aug. 2013).

59. Berisa, T. & Pickrell, J. K. Approximately independent linkage disequilibrium blocks in human populations. Bioinformatics 32, 283–285. issn: 1367-4803. 10.1093/bioinformatics/btv546 (2024) (Jan. 2016).

