## Supplemental Material for "Sparse modeling of interactions enables fast detection of genome-wide epistasis in biobank-scale studies"

### Supplementary Material

**Table S1. While using a mask that induces localized sparsity, SME produces deflated type I error rates when synthetic traits are generated under the null model.** Synthetic traits were simulated with only additive effects using chromosome 1 from individuals of self-identified European ancestry in the UK Biobank. These data were then subsampled using sample sizes of 20k, 50k, 100k, and 300k individuals. A total of 100 causal additive variants were randomly selected for each trait and their effects were assumed to explain 40% of the phenotypic variance. Data were analyzed using SME under varying percentages of SNPs that are masked (0%, 95%, and 99%, respectively). Empirical size for the analyses used significance thresholds of $\alpha$ = 0.05, 0.01, and 0.001. Values in the parentheses are the standard deviations of the estimates. Results are based on 100 simulations per scenario.

| Method | Sample Size | $\alpha$=0.05 | $\alpha$=0.01 | $\alpha$=0.001 |
|---|---|---|---|---|
| **SME (0% masked)** | 20k | 0.0414 (0.0181) | 0.0052 (0.0081) | 0.0000 (0.0000) |
| | 50k | 0.0427 (0.0199) | 0.0073 (0.0095) | 0.0005 (0.0022) |
| | 100k | 0.0474 (0.0234) | 0.0073 (0.0081) | 0.0006 (0.0024) |
| | 300k | 0.0614 (0.0279) | 0.0082 (0.0099) | 0.0011 (0.0032) |
| **SME (95% masked)** | 20k | 0.0204 (0.0148) | 0.0009 (0.0029) | 0.0000 (0.0000) |
| | 50k | 0.0232 (0.0178) | 0.0007 (0.0026) | 0.0000 (0.0000) |
| | 100k | 0.0278 (0.0164) | 0.0021 (0.0043) | 0.0000 (0.0000) |
| | 300k | 0.0359 (0.0201) | 0.0030 (0.0055) | 0.0005 (0.0021) |
| **SME (99% masked)** | 20k | 0.0044 (0.0066) | 0.0000 (0.0000) | 0.0000 (0.0000) |
| | 50k | 0.0039 (0.0069) | 0.0000 (0.0000) | 0.0000 (0.0000) |
| | 100k | 0.0048 (0.0069) | 0.0000 (0.0000) | 0.0000 (0.0000) |
| | 300k | 0.0077 (0.0080) | 0.0002 (0.0015) | 0.0000 (0.0000) |

**Table S2. Complete list of marginal epistasis results from running SME on mean corpuscular hemoglobin (MCH) assayed in individuals in the UK Biobank.** Here, we analyze 349,411 white British individuals in the UK Biobank genotyped at 543,813 SNPs genome-wide. As a mask, we leveraged DNase I-hypersensitive sites (DHS) data measured over 12 days of *ex vivo* erythroid differentiation [28, 34].In the first three columns, we list the identifier of the SNPs, their chromosome, and their genomic location (basepair position). Next, we give the $P$-value and marginal epistatic phenotypic variance explained (PVE) for each SNP as esimtated by SME. In the last two columns, we give the abbreviation for the trait that is analyzed and the type of the external data source used to induce sparsity. These results are saved as comma-separated value (CSV) files and can be downloaded directly from https://doi.org/10.5281/zenodo.14607998.

**Table S3. Complete list of marginal epistasis results from running SME on mean corpuscular hemoglobin concentration (MCHC) assayed in individuals in the UK Biobank.** Here, we analyze 349,411 white British individuals in the UK Biobank genotyped at 543,813 SNPs genome-wide. As a mask, we leveraged DNase I-hypersensitive sites (DHS) data measured over 12 days of *ex vivo* erythroid differentiation [28, 34]. In the first three columns, we list the identifier of the SNPs, their chromosome, and their genomic location (basepair position). Next, we give the $P$-value and marginal epistatic phenotypic variance explained (PVE) for each SNP as esimtated by SME. In the last two columns, we give the abbreviation for the trait that is analyzed and the type of the external data source used to induce sparsity. These results are saved as comma-separated value (CSV) files and can be downloaded directly from https://doi.org/10.5281/zenodo.14607998.

**Table S4. Complete list of marginal epistasis results from running SME on mean corpuscular volume (MCV) assayed in individuals in the UK Biobank.** Here, we analyze 349,411 white British individuals in the UK Biobank genotyped at 543,813 SNPs genome-wide. As a mask, we leveraged DNase I-hypersensitive sites (DHS) data measured over 12 days of *ex vivo* erythroid differentiation [28, 34]. In the first three columns, we list the identifier of the SNPs, their chromosome, and their genomic location (basepair position). Next, we give the $P$-value and marginal epistatic phenotypic variance explained (PVE) for each SNP as esimtated by SME. In the last two columns, we give the abbreviation for the trait that is analyzed and the type of the external data source used to induce sparsity. These results are saved as comma-separated value (CSV) files and can be downloaded directly from https://doi.org/10.5281/zenodo.14607998.

**Table S5. Complete list of marginal epistasis results from running SME on hematocrit (HCT) assayed in individuals in the UK Biobank.** Here, we analyze 349,411 white British individuals in the UK Biobank genotyped at 543,813 SNPs genome-wide. As a mask, we leveraged DNase I-hypersensitive sites (DHS) data measured over 12 days of *ex vivo* erythroid differentiation [28, 34]. In the first three columns, we list the identifier of the SNPs, their chromosome, and their genomic location (basepair position). Next, we give the $P$-value and marginal epistatic phenotypic variance explained (PVE) for each SNP as esimtated by SME. In the last two columns, we give the abbreviation for the trait that is analyzed and the type of the external data source used to induce sparsity. These results are saved as comma-separated value (CSV) files and can be downloaded directly from https://doi.org/10.5281/zenodo.14607998.

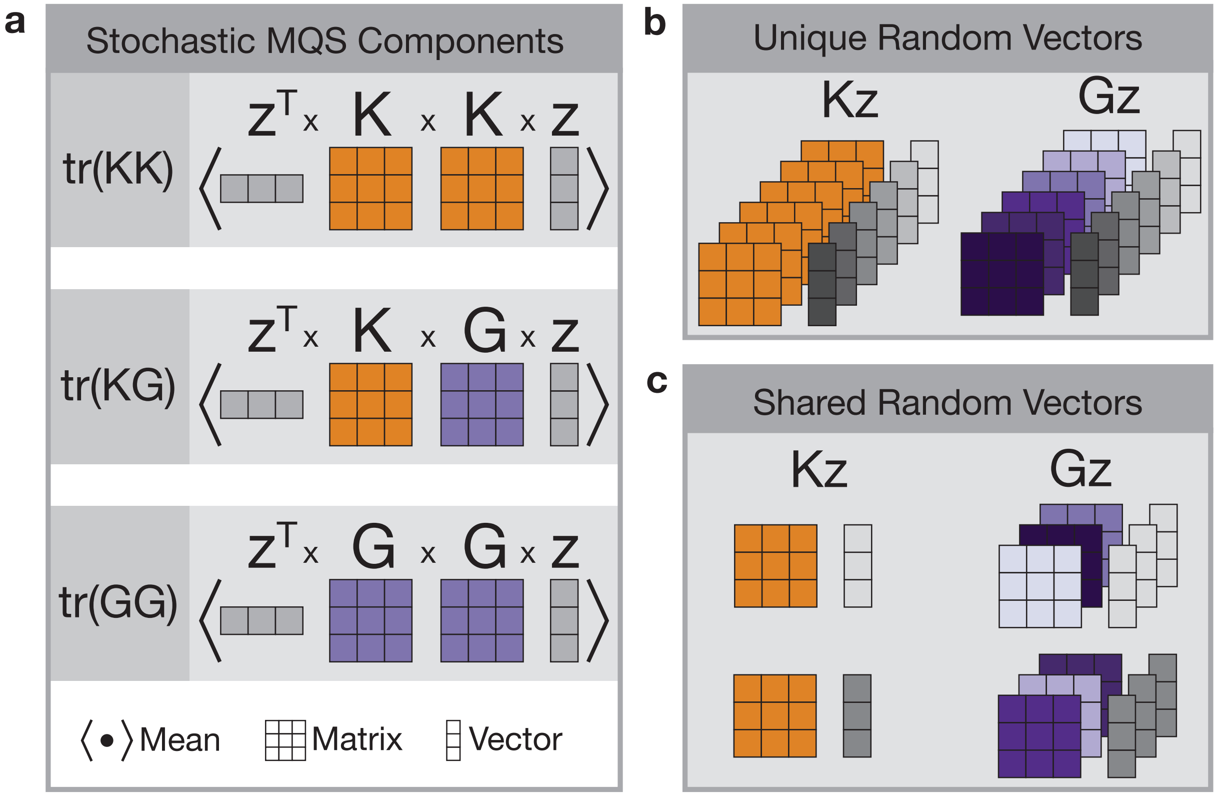


**Figure S1. Schematic overview illustrating the novel approximation to the stochastic trace estimator used in SME. (a)** Computing the exact trace of a product for two covariance matrices is computationally infeasible for large studies with many individuals. SME overcomes this limitation by computing point estimates for variance components using a method-of-moments (MoM) algorithm that features traces of matrix products with random vectors $\mathbf{z}$. In the stochastic trace estimates, we can identify reusable matrix-by-vector products. We see that the matrix-by-vector products of the form $\mathbf{Az}$ with $\mathbf{A} \in \{\mathbf{K}, \mathbf{G}_j\}$ and combinations thereof appear in multiple traces. **(b)** The genetic relatedness matrix $\mathbf{K}$ is the same for all focal SNPs. Using unique random vectors in this computation for every focal SNP, we compute the stochastic approximation repeatedly. Computing the matrix-by-vector products $\mathbf{Kz}$ constitutes about half of the computation time of the point estimates when no mask is applied. It constitutes most of the computation time when a mask induces sparsity. **(c)** By sharing random vectors $\mathbf{z}$ between a set of focal SNPs, computing $\mathbf{Kz}$ can be done once for the entire set. With this, we greatly reduce the computational burden of computing $\mathbf{Kz}$.

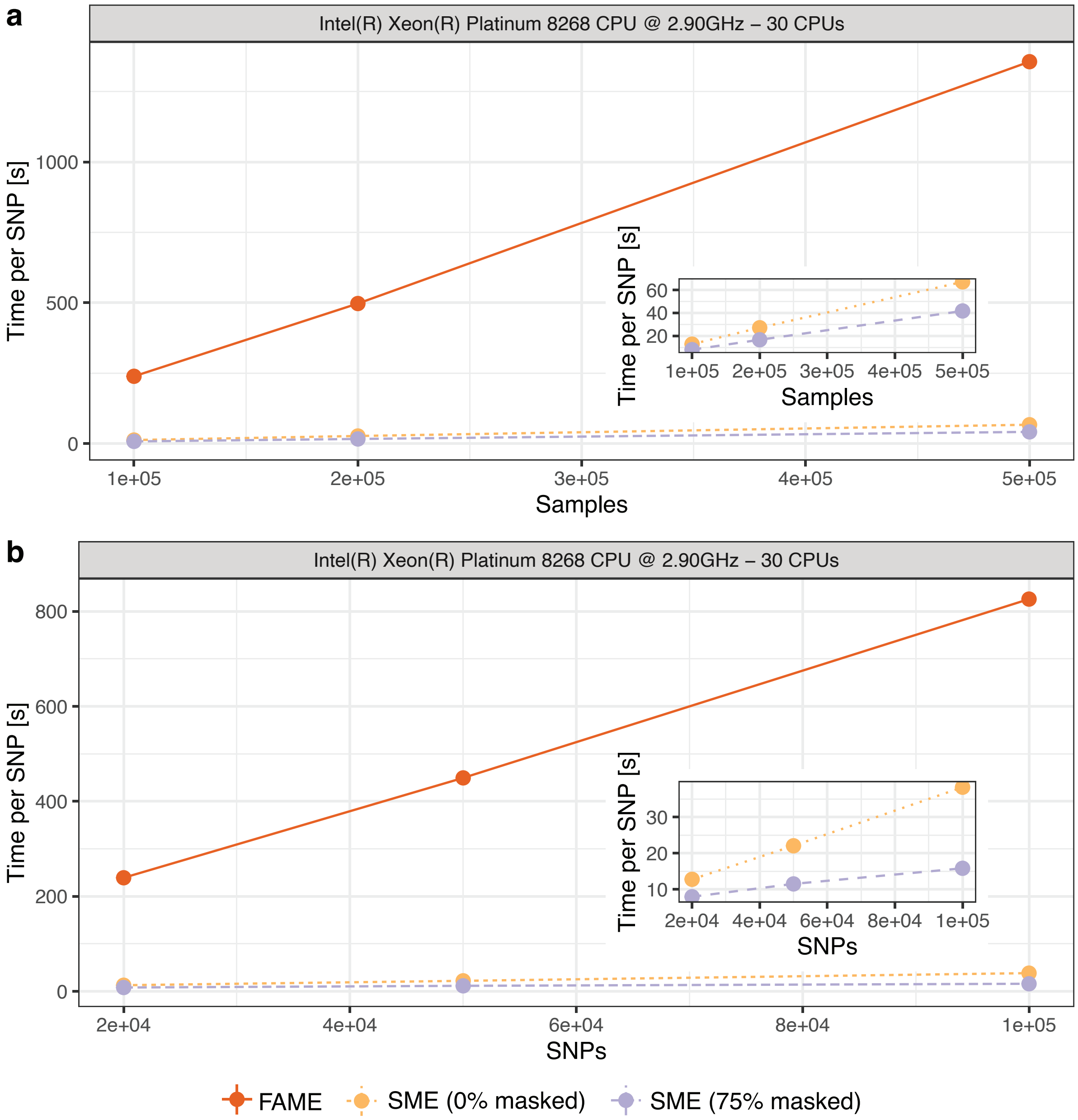


**Figure S2. Computational time needed to run a single test with SME and FAME [4] as a function of sample size and total number of SNPs.** The solid orange line represents the runtime using FAME. The dotted yellow line shows the runtime of SME without a mask. The dashed purple line represents the runtime of SME with 75% of variants masked. In all cases, the number of random vectors is fixed at 100. The number of variants sharing random vectors in SME is fixed at 90. Computations were performed on an Intel(R) Xeon(R) Platinum 8268 CPU with 30 cores. **(a)** Computational time per test (in seconds) as a function of sample size, with the genome size fixed at 20,000 SNPs. **(b)** Computational time per test (in seconds) as a function of the genome size in number of SNPs, with the sample size fixed at 100,000 individuals.

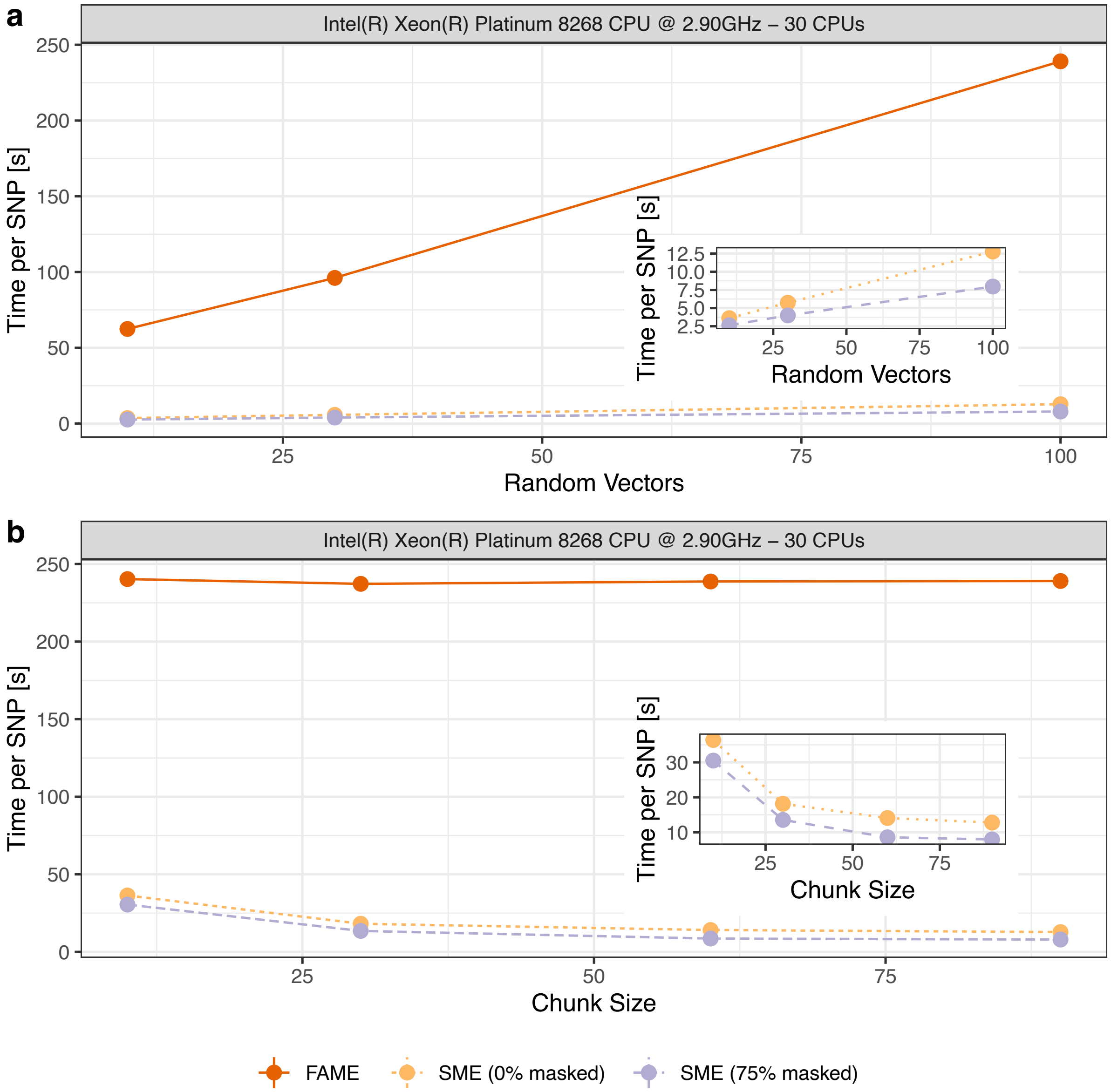


**Figure S3. Computational time needed to run a single test with SME and FAME [4] as a function of the number of random vectors and number of SNPs sharing random vectors.** The solid orange line represents the runtime using the command-line tool FAME. The dotted yellow line shows the runtime using the `R` function of SME without applying a mask. The dashed purple line represents the runtime using SME with 75% of the variants masked. The analyzed data had a genome size of 20,000 SNPs and a sample size of 100,000 individuals. All computations were performed on an Intel(R) Xeon(R) Platinum 8268 CPU with 30 cores. **(a)** Computational time per test as a function of the number of random vectors used in the stochastic trace estimator. The number of focal variants sharing random vectors (chunk size) in SME was fixed at 90. **(b)** Computational time per test as a function of the number of focal variants sharing random vectors (chunk size). The number of random vectors was fixed at 100.

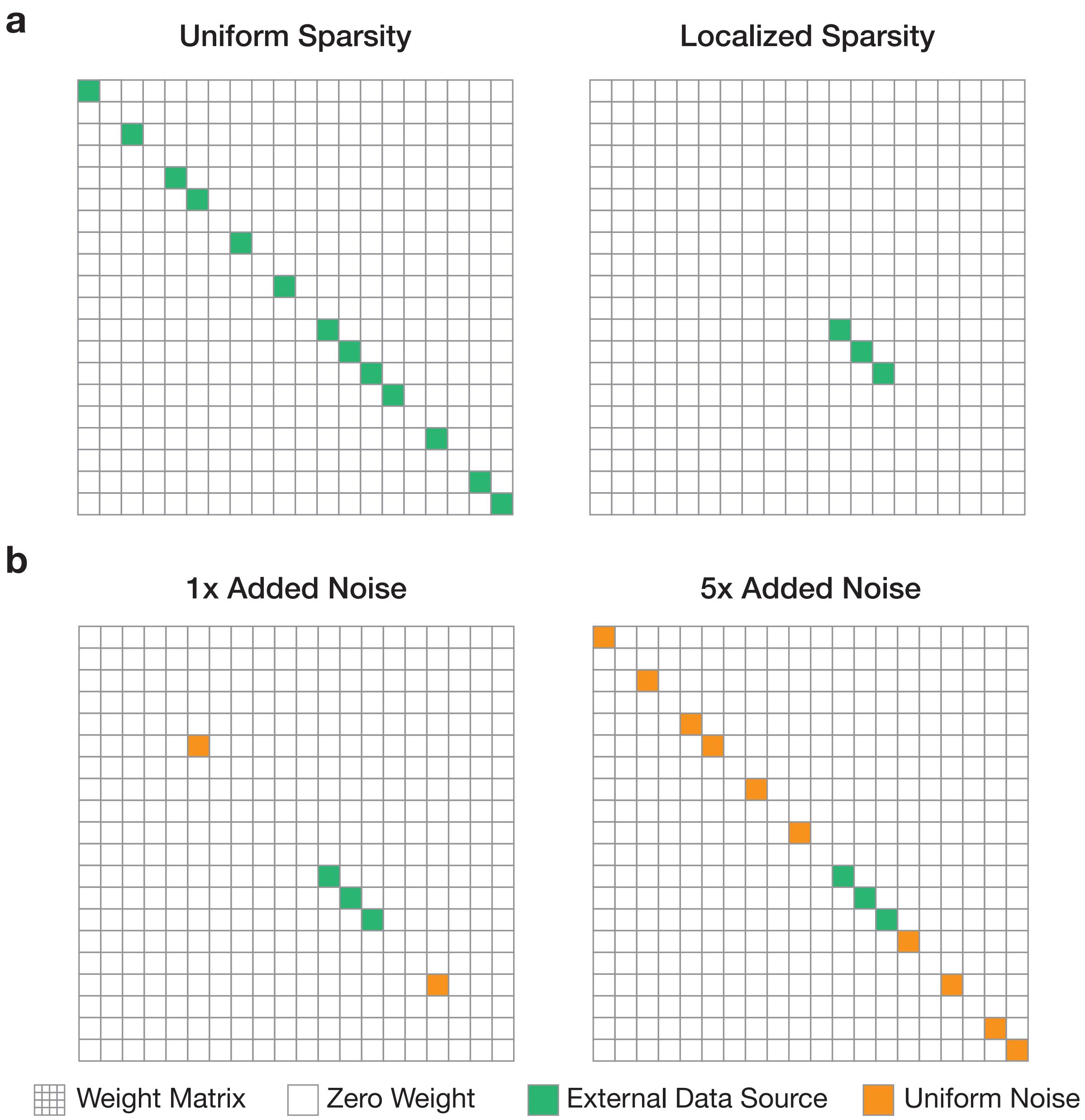


**Figure S4. Schematic overview illustrating weight matrices with uniform and localized sparsity.** **(a)** The simulations consider two scenarios that result in different distributions of sparsely modeled gene-interactions. The weight matrices are binary everywhere except for places on the diagonal where variants satisfy some criteria from an external data source (e.g., it is located in a position on the genome that overlaps with open chromatin regions). The first column illustrates *uniform sparsity* where non-zero entries are evenly distributed along the diagonal. As a result the modeled gene-interactions are evenly distributed along the genome. In the second scenario, the external data source induces *localized sparsity* where the modeled gene-interactions are concentrated in a small genomic window illustrated as a block structure in the weight matrix. **(b)** SME produces negatively biased variance component estimates when using an external data source that induces localized sparsity in the model. To overcome this issue in practice, we propose a strategy in which we take an external data source with localized genomic information and randomly unmask "unimportant" variants with uniform probability along the genome (making the localized sparsity look more uniform). The cartoon illustrates the distribution of non-zero weights in the weight matrix after including an additional 1× and 5× of initially disregarded interactions.

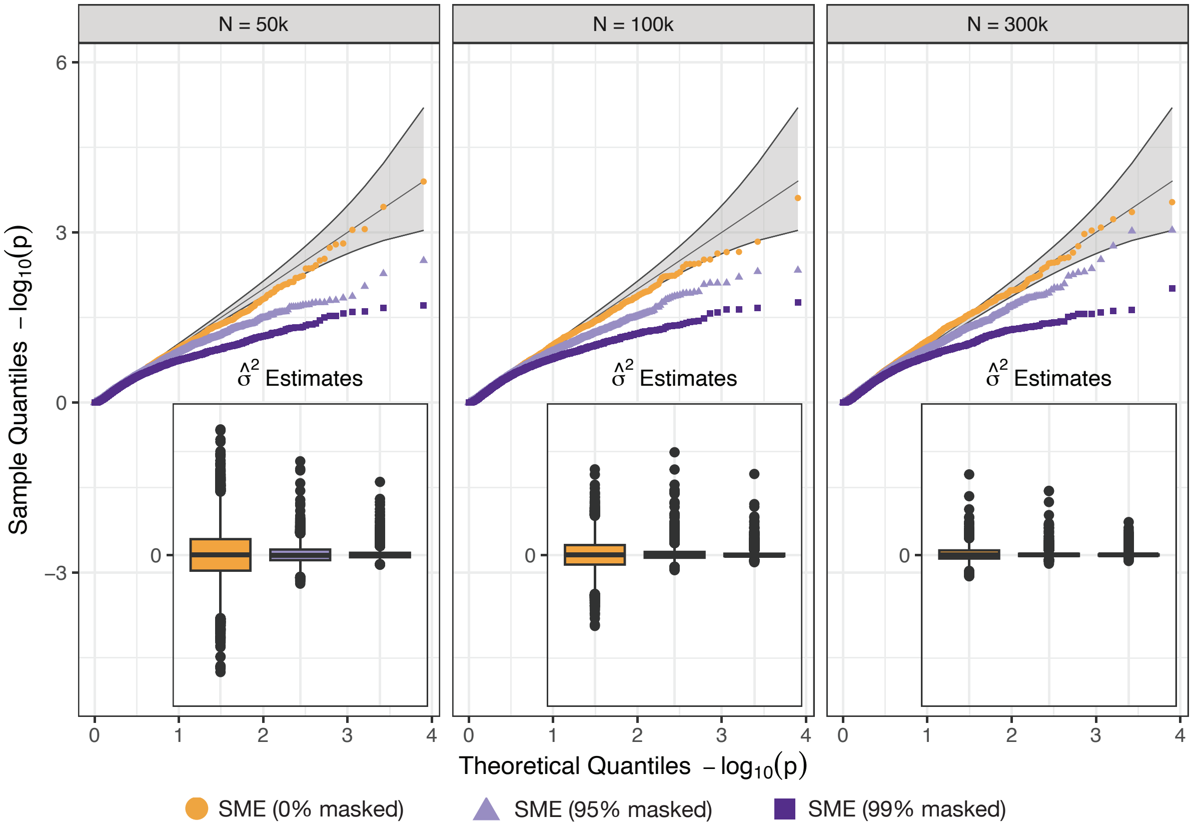


**Figure S5. While using a mask that induces localized sparsity, SME is negatively biased under the null hypothesis.** Synthetic traits were simulated with only additive effects using chromosome 1 from individuals of self-identified European ancestry in the UK Biobank. These data were then subsampled using sample sizes of 50k, 100k, and 300k individuals. A total of 100 causal additive variants were randomly selected for each trait and their effects were assumed to explain 40% of the phenotypic variance. To simulate localized sparsity in the mask, we randomly sample a seed SNP and define a block around it. All SNPs not in that block are masked. Data were analyzed with SME under varying percentages of SNPs that are masked (0%, 95%, and 99%, respectively). The small insets in each plot show the distribution of the estimated marginal epistatic variance components across all experiments. For reference, under the null hypothesis $H_0 : \sigma^2 \leq 0$. Results are based on 100 simulations per scenario.

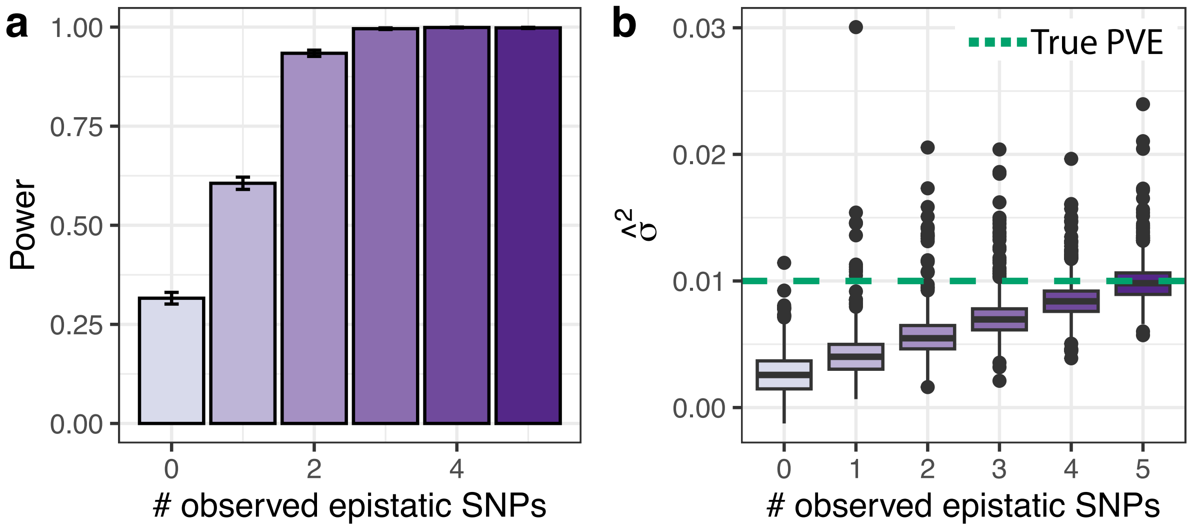


**Figure S6. Consequence of having a misspecified mask on power and variance component estimates in SME.** Synthetic traits were simulated using chromosome 1 from individuals of self-identified European ancestry in the UK Biobank. Data were subsampled to 100k individuals. We randomly selected 10% of all variants to have additive effects that collectively explained 30% of the trait variance. We then fixed the total epistatic variance to 5%. Each epistatic SNP was simulated to have 5 interaction partners. SME was given a weight matrix in which 95% of all SNPs were masked. In this simulation, we investigate a scenario where the weight matrix incorrectly masked 0 to 5 of true interacting partners for each causal SNP. **(a)** Empirical power computed as the function of masked true interactive partners. Here, significant SNPs were evaluated at a genome-wide threshold $P < 5 \times 10^{-8}$. Error bars represent the standard deviation across 100 replicate experiments. **(b)** Marginal epistatic variance component estimates ($\hat{\sigma}^2$) as a function of masked true interactive partners. The dashed green line represents the true simulated per SNP epistatic phenotypic variance explained (PVE).

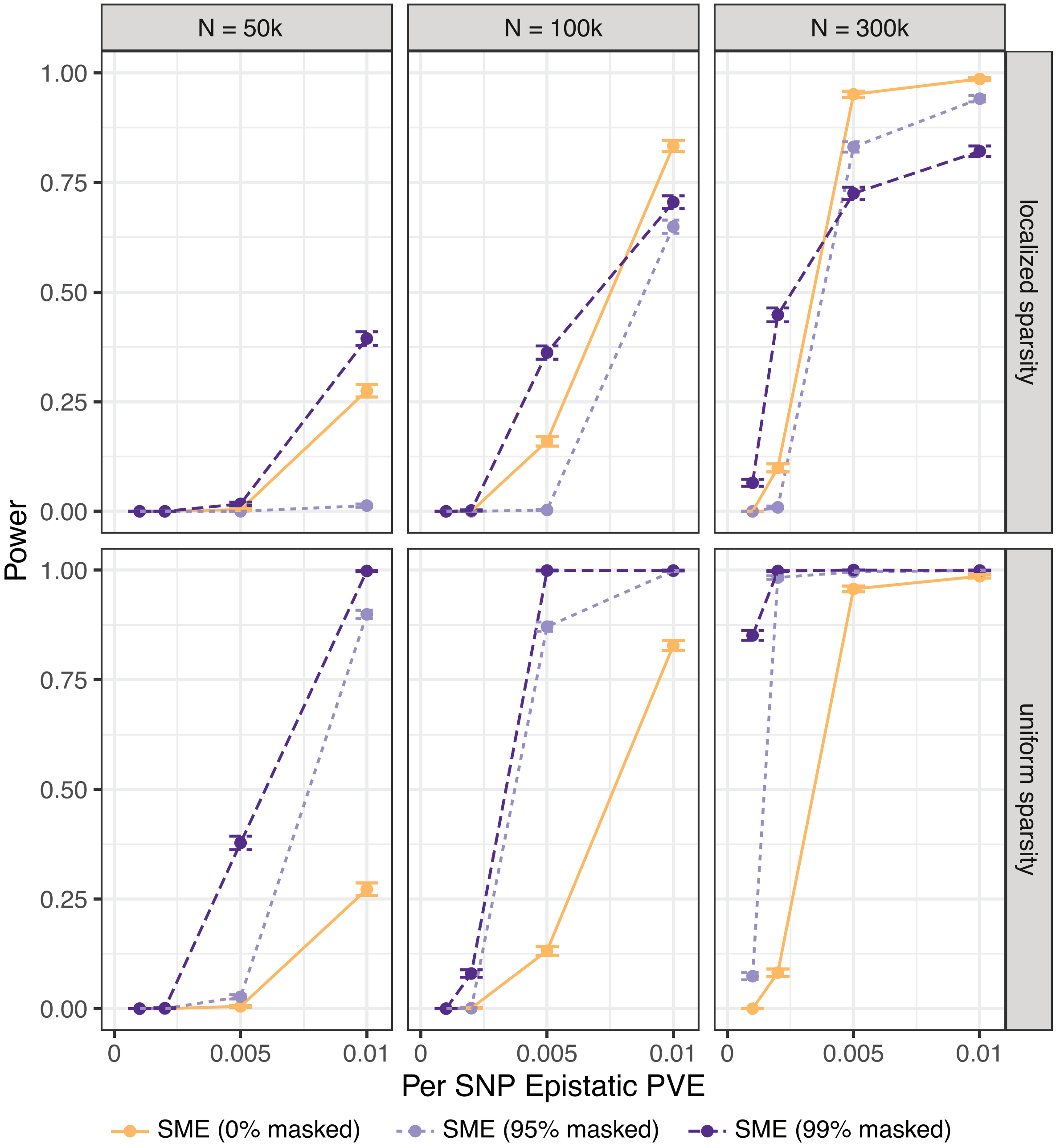


**Figure S7. SME exhibits less power when using a mask that induces localized sparsity.** Synthetic traits were simulated with both additive and pairwise epistatic effects using chromosome 1 from individuals of self-identified European ancestry in the UK Biobank. Data were subsampled using sample sizes of 50k, 100k, and 300k individuals. We randomly selected 10% of all variants to have additive effects that collectively explained 30% of the trait variance. We then fixed the total epistatic variance to 5%. The per SNP epistatic phenotypic variance explained (PVE) was adjusted by varying the number of interacting SNPs (chosen to be 10, 20, 50, or 100 SNPs). Data were analyzed using SME under varying percentages of variants that are excluded from consideration as potential interaction partners for each focal SNP (0%, 95%, and 99% masking, respectively). Empirical power was determined using the significance threshold $P < 5 \times 10^{-8}$. Results for uniform sparsity are given on the bottom as reference. We conducted 100 simulations per scenario, with error bars representing the standard deviation across replicates.

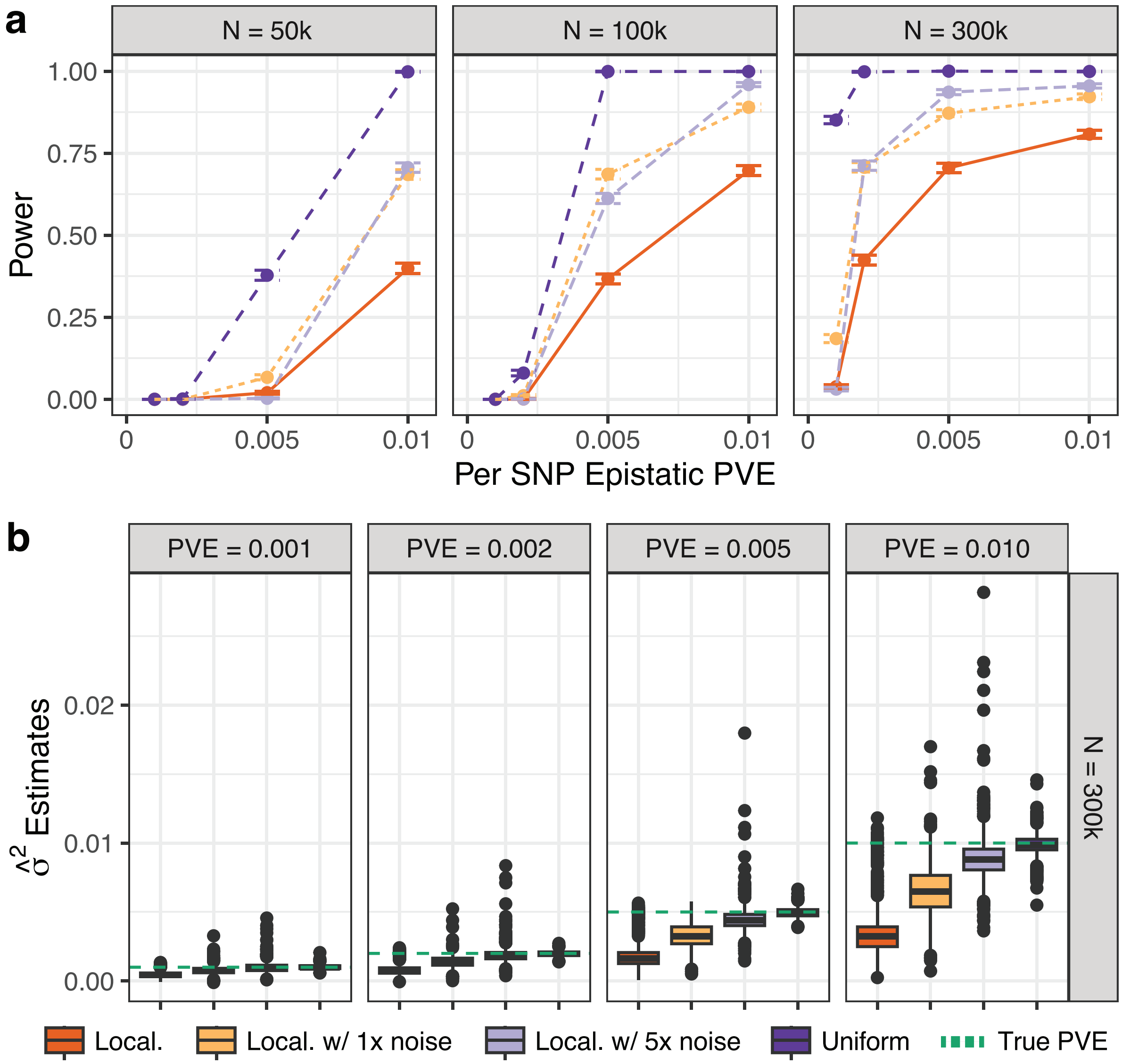


**Figure S8. Adding random noise to external data sources that induce localized sparsity recoveres empirical power and reduces the negative biase in variance component estimates.** Synthetic traits were simulated with both additive and pairwise epistatic effects using chromosome 1 from individuals of self-identified European ancestry in the UK Biobank. Data were subsampled using sample sizes of 50k, 100k, and 300k individuals. We randomly selected 10% of all variants to have additive effects that collectively explained 30% of the trait variance. We then fixed the total epistatic variance to 5%. The per SNP epistatic phenotypic variance explained (PVE) was adjusted by varying the number of interacting SNPs (chosen to be 10, 20, 50, or 100 SNPs). We simulated external data sources that induced both *uniform sparsity* (dark purple) and *localized sparsity* (dark orange) resulting in 99% of variants being masked. To counteract the negative bias of the localized sparsity, we randomly unmask 1% and 5% additional SNPs with uniform probability along the entire chromosome—making the localized sparsity look more uniform (also see Fig. S4). **(a)** Empirical power at a genome-wide significance threshold $P < 5 \times 10^{-8}$, shown as a function of the per SNP epistatic phenotypic variance explained (PVE) by causal variants across three different sample sizes. Error bars represent the standard deviation across 100 replicate experiments. **(b)** Marginal epistatic variance component estimates ($\hat{\sigma}^2$) on simulated traits with 300k individuals. The dashed green line represents the true per SNP epistatic PVE.

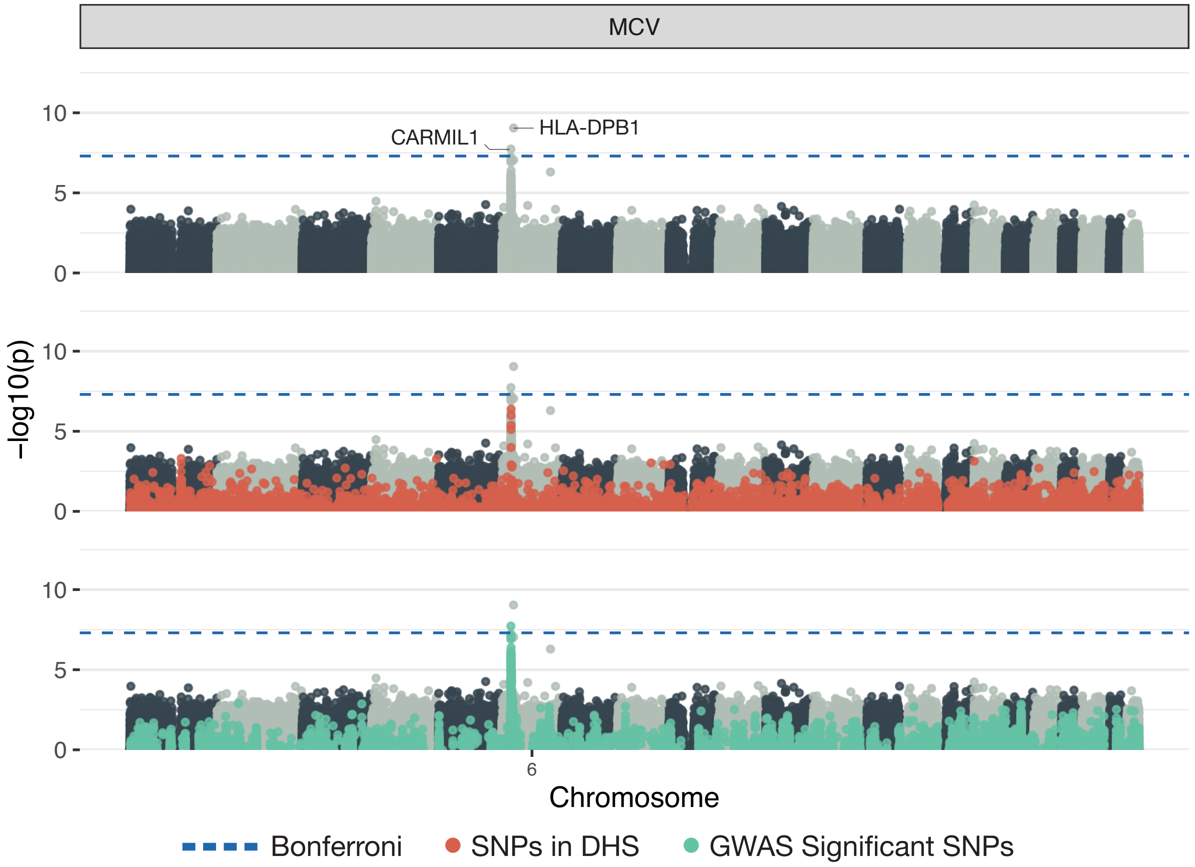


**Figure S9. Manhattan plots of a genome-wide interaction analysis using SME to study mean corpuscular volume (MCV) assayed in individuals in the UK Biobank.** As a mask in this study, we leveraged DNase I-hypersensitive sites (DHS) data measured over 12 days of *ex vivo* erythroid differentiation [28, 34]. This means that, while all SNPs are tested for marginal epistasis, only their interactions with SNPs in DHS regions are considered. Here, $-\log_{10}$ transformed $P$-values from SME are plotted for each SNP against their genomic positions. Chromosomes are shown in alternating colors for clarity. The dashed blue line represents the genome-wide significance threshold ($P < 5 \times 10^{-8}$). Each panel shows the same plot with different aspects of the result highlighted. The first simply shows the names of the closest neighboring genes to significant epistatic SNPs. The second panel highlights the SNPs that fall in DHS regions, and the third panel highlights SNPs that were also found to have a significant (additive) association with the trait according to a GWAS.

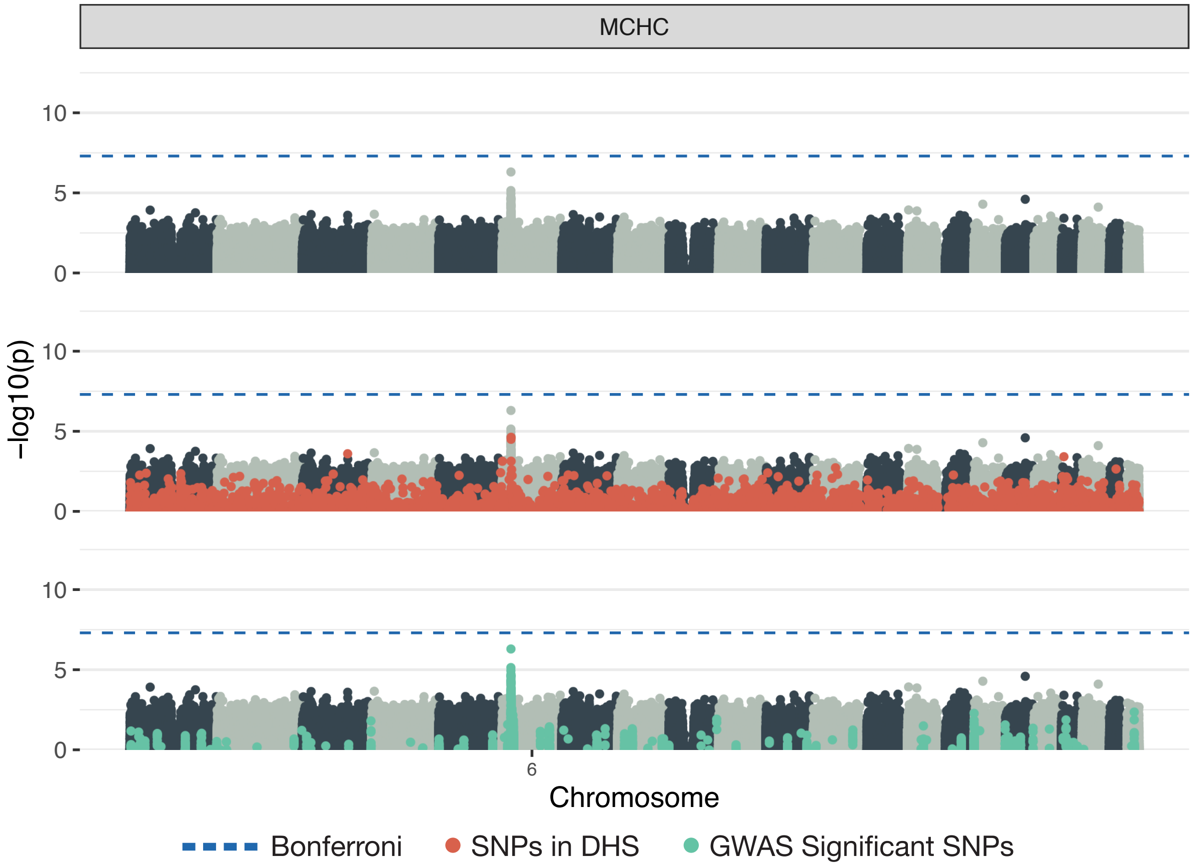


**Figure S10. Manhattan plots of a genome-wide interaction analysis using SME to study mean corpuscular hemoglobin concentration (MCHC) assayed in individuals in the UK Biobank.** As a mask in this study, we leveraged DNase I-hypersensitive sites (DHS) data measured over 12 days of *ex vivo* erythroid differentiation [28, 34]. This means that, while all SNPs are tested for marginal epistasis, only their interactions with SNPs in DHS regions are considered. Here, $-\log_{10}$ transformed $P$-values from SME are plotted for each SNP against their genomic positions. Chromosomes are shown in alternating colors for clarity. The dashed blue line represents the genome-wide significance threshold ($P < 5 \times 10^{-8}$). Each panel shows the same plot with different aspects of the result highlighted. The first simply shows the names of the closest neighboring genes to significant epistatic SNPs. The second panel highlights the SNPs that fall in DHS regions, and the third panel highlights SNPs that were also found to have a significant (additive) association with the trait according to a GWAS.

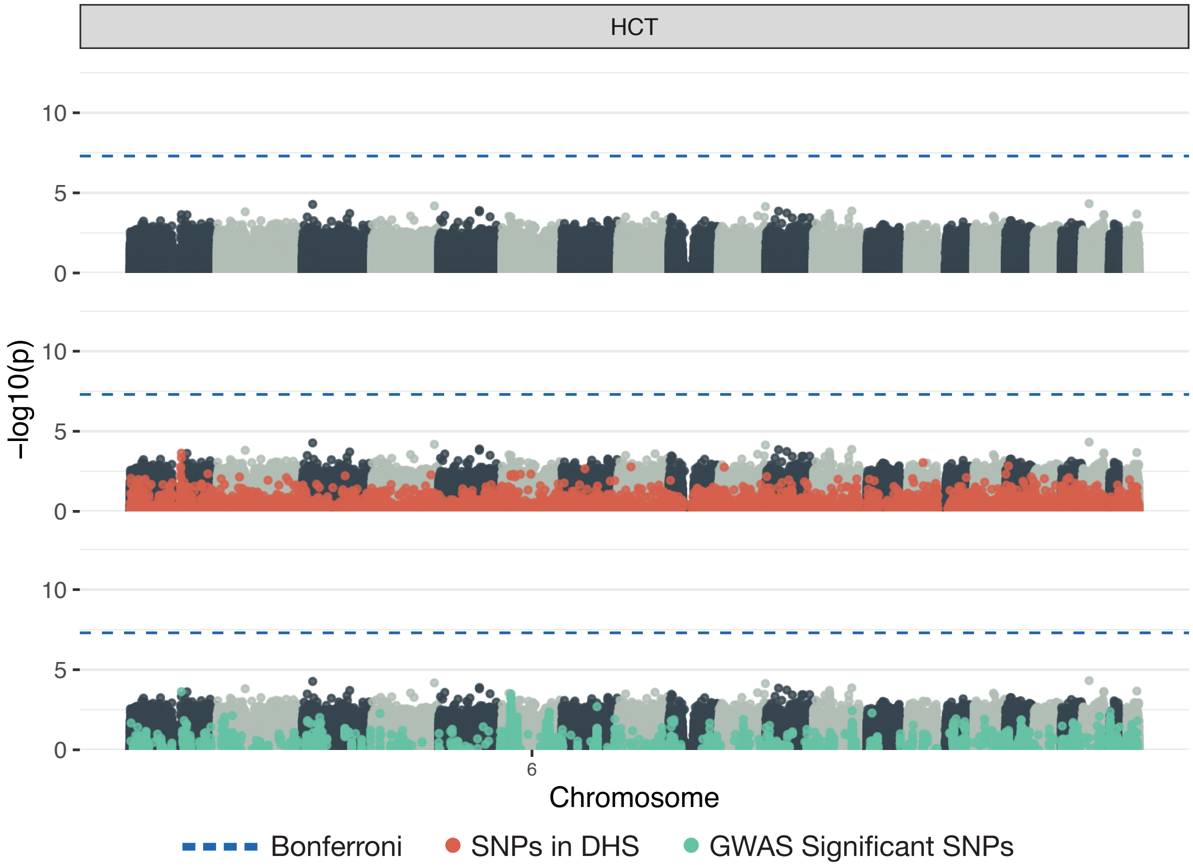


**Figure S11. Manhattan plots of a genome-wide interaction analysis using SME to study hematocrit (HCT) assayed in individuals in the UK Biobank.** As a mask in this study, we leveraged DNase I-hypersensitive sites (DHS) data measured over 12 days of *ex vivo* erythroid differentiation [28, 34]. This means that, while all SNPs are tested for marginal epistasis, only their interactions with SNPs in DHS regions are considered. Here, $-\log_{10}$ transformed $P$-values from SME are plotted for each SNP against their genomic positions. Chromosomes are shown in alternating colors for clarity. The dashed blue line represents the genome-wide significance threshold ($P < 5 \times 10^{-8}$). Each panel shows the same plot with different aspects of the result highlighted. The first simply shows the names of the closest neighboring genes to significant epistatic SNPs. The second panel highlights the SNPs that fall in DHS regions, and the third panel highlights SNPs that were also found to have a significant (additive) association with the trait according to a GWAS.

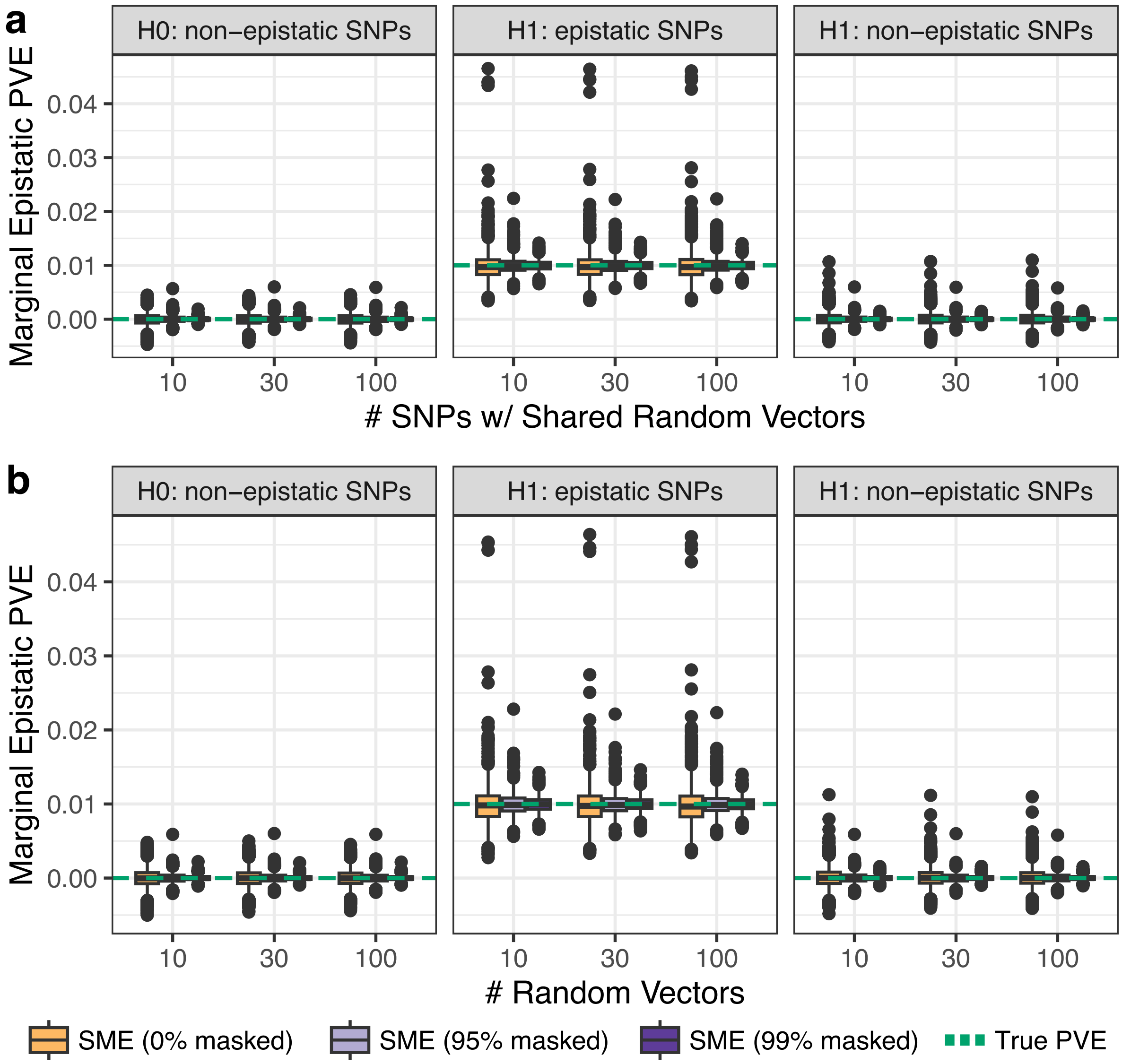


**Figure S12. The SME stochastic method-of-moments algorithm produces estimates that are robust to the choice of number of random vectors and the number of SNPs sharing random vectors.** Synthetic traits were simulated under the null hypothesis ($H_0$) of no epistasis and the alternaitve hypothesis ($H_1$) using chromosome 1 from individuals of self-identified European ancestry in the UK Biobank 100k individuals. We randomly selected 10% of all variants to have additive effects that collectively explained 30% of the trait variance. For simulations under $H_1$, we then fixed the total epistatic variance to 5%. The per SNP epistatic phenotypic variance explained (PVE) was adjusted by randomly choosing 10 epistatic SNPs. **(a)** Marginal epistatic variance component estimates as a function of the number of SNPs sharing random vectors in the stochastic trace estimates (also see Fig. S1). The number of random vectors was fixed at 100. **(b)** Marginal epistatic variance component estimates as a function of number of random vectors in the stochastic trace estimates. The number of SNPs sharing random vectors was fixed at 100. In both panels, the orange box plots show the results of using SME with no masking applied. The light and dark purple show the results for implementing SME with 95% and 99% masking, respectively. The dashed green line indicates the true marginal epistatic PVE used in the simulations.
